# Liquid condensates increase potency of amyloid fibril inhibitors

**DOI:** 10.1101/2020.10.29.360206

**Authors:** Thomas C. T. Michaels, L. Mahadevan, Christoph A. Weber

## Abstract

In living cells, liquid condensates form in the cytoplasm and nucleoplasm via phase separation and regulate physiological processes. They also regulate aberrant aggregation of amyloid fibrils, a process linked to Alzheimer’s and Parkinson’s diseases. In the absence of condensates it has been shown that amyloid aggregation can be inhibited by molecular chaperones and rationally designed drugs. However it remains unknown how this drug- or chaperone-mediated inhibition of amyloid fibril aggregation is affected by phase-separated condensates. Here we study the interplay between protein aggregation, its inhibition and liquid-liquid phase separation. Our key finding is that the potency of inhibitors of amyloid formation can be strongly enhanced. We show that the corresponding mechanism relies on the colocalization of inhibitors and aggregates inside the liquid condensate. We provide experimentally testable physicochemical conditions under which the increase of inhibitor potency is most pronounced. Our work highlights the role of spatio-temporal heterogeneity in curtailing aberrant protein aggregation and suggests design principles for amyloid inhibitors accounting for partitioning of drugs into liquid condensates.

---

The proteostasis network in living cells uses chaperones as key players for protein quality control [1, 2]. The main role of chaperones is to stabilize the native fold of proteins but they are also involved in degradation processes such as lysosome-mediated degradation of dysfunctional proteins [3, 4]. A decline of the proteostasis network, due to ageing for example [2], causes the accumulation of misfolded proteins and their subsequent aggregation into amyloid fibrils – a process linked to over 50 devastating disorders, including Alzheimer’s and Parkinson’s diseases [5–9]. It is therefore important to obtain insights into the mechanisms underlying the aggregation of misfolded proteins into amyloid fibrils.

These mechanistic insights are particularly important for understanding how inhibitors affect amyloid fibril formation. A range of compounds, including molecular chaperones [10] or rationally designed inhibitors, such as small drug-like molecules [11, 12], nanoparticles [13], and antibodies [14], have shown to affect amyloid formation *in vitro* and *in vivo* [11–27]. Recent *in vitro* studies indicate that these compounds inhibit aggregation of amyloid fibrils by binding to specific of targets, including monomers, fibril ends or surfaces [10, 28–31]. However, these different inhibition mechanisms have been elucidated in homogeneous *in vitro* systems, disregarding that cells are spatially heterogeneous due to the presence of organelles.

A new class of organelles that has recently received significant attention are protein condensates. They form via phase separation from the cyto- or nucleoplasm and share most physical properties with liquid-like, condensed droplets [32–35]. Liquid condensates provide physiochemical environments distinct from the surrounding cyto- and nucleoplasm, allowing to spatially regulate a range of biochemical reactions associated with biological function or dysfunction. Recent experimental [35–37] and theoretical [38] studies show that condensates can affect amyloid aggregation. For example, stress granules concentrate monomers causing aggregation to occur only inside the condensates [39, 40]. Condensates can accumulate chaperones capable of either suppressing aberrant phase transition of condensates [41] or driving amyloid formation by sequestering misfolded proteins [42–44]. Chaperones are also involved in the control of condensate stability [45] and the surveillance of stress granule quality [46, 47].

The relevance of liquid condensates for the spatial organization of chaperones and amyloids suggests that spatial compartmentalisation may play an important role in regulating and suppressing aberrant protein aggregation. Similarly, the action of synthetic drugs capable of inhibiting amyloid formation may be affected by the presence of liquid condensates. Abbreviating both chaperones and synthetic inhibitors as ‘drugs’, we raise the following question: how do liquid condensates affect drug-mediated inhibition of amyloid fibril formation (Fig. 1)? Here, we address this question by presenting a physical model of amyloid inhibition coupled to liquid-liquid phase separation. Our model is inspired by recent experimental studies on amyloid formation in the presence of condensates and builds on the latest insights obtained from systematic studies on homogeneous amyloid aggregation kinetics in the presence of inhibiting drugs. The key phase separation parameters determining the amplification are the condensate volume and the partitioning coefficients of drug and monomers. Using our model we find that liquid condensates can significantly amplify drug-mediated inhibition of amyloid fibrils. The mechanism for this amplification builds on the colocalization inside the condensate of the inhibiting drug and the target site such as the aggregating monomers or the aggregates themselves (Fig. 1(c-d)). Our results have important implications for the control of aggregation in cells and for drug discovery against amyloid disorders. In fact, our work showcases the potential of condensed phases to regulate aberrant protein aggregation in living cells. Moreover, our findings suggest new design principles for amyloid inhibitors based on pronounced partitioning of drugs into the liquid condensates.

**FIG. 1.**
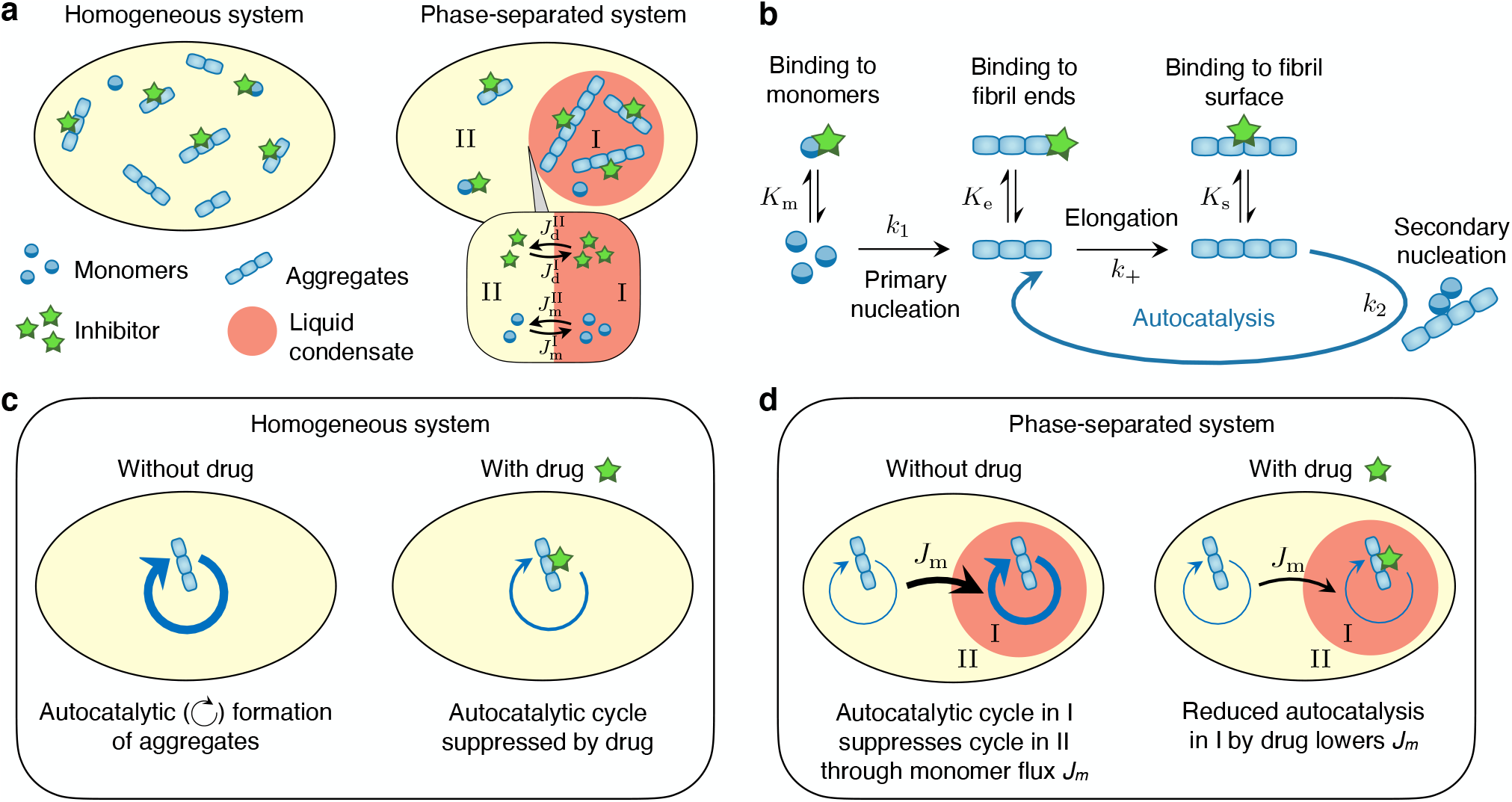
Impact of liquid-like condensates on drug-mediated inhibition of amyloid aggregation. **(a)** We determine the physical conditions when the presence of a phase-separated condensate (phase I, right) yields a higher degree of inhibition (e.g., less fibrils at the end of the aggregation reaction) compared to a homogeneous system (left). Inset: Definition of partitioning of monomers and drug components and associated non-equilibrium diffusive fluxes across the condensate interface. We neglect diffusive exchange of fibrils through the interface because long fibrils may only slowly diffuse due to the presence of other fibrils (repatation). **(b)** The reaction network of amyloid fibril assembly and three established mechanisms of inhibition [10, 28–30]. **(c-d)** Mechanism by which an inhibitor of secondary nucleation affects amyloid aggregation in a homogeneous **(c)** and a heterogeneous system **(d)**. In a homogeneous system the drug reduces the overall fibril concentration by suppressing the autocatalytic fibril cycle associated with secondary nucleation and growth (depicted by a circular arrow, see panel (b)). In the phase-separated system, partitioning the drug in I suppresses the autocatalytic fibril cycle in the condensate. This causes a decrease of 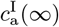, but also a reduction of the monomer flux *Jm* from II to I, which in turn causes 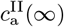 to rise. The balance between these two opposing effects determines the efficacy of the drug to suppress aggregation in the heterogeneous setting.

## Model for amyloid formation with inhibitors and liquid condensates

As the starting point of our model of amyloid inhibition with liquid condensates, we consider a system of volume *V* containing a phase-separated liquid-like condensate of volume *V* ^I^ (phase I) coexisting with the surrounding, dilute phase (phase II). Phase I is rich in protein *A*, while the other phase is dilute in *A* but could be rich in another protein, lipids or water, shortly denoted as *B* component (Fig. 1(a)). In addition to the phase-separating components *A*, *B*, the system contains a concentration 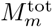 of amyloidogenic monomers and a concentration *c*_d_ of an inhibitor of amyloid aggregation. We assume that all aggregating species and the drug are sufficiently dilute with respect to *A* and *B* such that their impact on *AB*-phase separation can be neglected (SI Sec. 1.3.1). The coexisting phases I and II thus create distinct biochemical environments for aggregation and give rise to diffusive fluxes across the interface that partition aggregating monomers and drug components, which we denote with *i* (i.e. *i* denotes monomers or inhibitors). By equating chemical potentials of component *i* between the two phases, we can calculate the equilibrium partitioning coefficient of *i*, 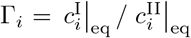 where 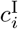 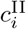 are the respective concentrations of component *i* in the phase I and II. Γ_*i*_ is determined by the degree of phase separation and the relative strength of interaction between species *i* and *A* or *B* (SI Sec. 1.3.1 and Ref. [38]). Favouring interactions between *i* and *A*, for instance, lead to Γ_*i*_ > 1, i.e., enrichment of *i* in the condensate. Deviations from partitioning equilibrium lead to inter-phase fluxes of species *i* that re-establish equilibrium. For small deviations to equilibrium, these fluxes depend linearly on the concentrations in phase I and II (SI Sec. S1.3.2 and Ref. [38])

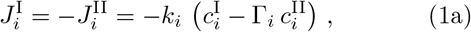

where Γ_*i*_ is the equilibrium partitioning coefficient and *k*_*i*_ = 4*πRD*_*i*_ is the relaxation rate toward equilibrium, with *R* being the radius of the condensate and *D*_*i*_ the diffusion coefficient of species *i*. Note that these fluxes vanish at equilibrium, i.e. when 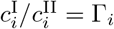.

In the presence of a phase-separated liquid condensate, aggregation of monomers into amyloid fibrils occurs in each phase I and II involving a number of different microscopic events [48–55], which are illustrated in Fig. 1(b). Aggregation is initiated by primary nucleation, i.e., the formation of the smallest fibrillar structures directly from the monomers. Aggregates then grow by fibril elongation. In addition, aggregation is often accelerated by secondary nucleation mechanisms [56], including fibril fragmentation, lateral branching or surface-catalyzed secondary nucleation (Fig. 1(b)). These secondary nucleation processes are characterised by a dependence on the current population of aggregates and introduce therefore an autocatalytic cycle in the system (Fig. 1(b)). We focus here on suppression of fibril kinetics by the inhibitor through three established molecular mechanisms [10, 28–30], including inhibitor binding to (i) free monomers, (ii) fibril ends, and (iii) fibril surface sites, as depicted in Fig. 1(b) (see SI Fig. S1 for examples of inhibitors of amyloid formation). For simplicity, we neglect drug binding to intermediate oligomers, but note that our framework can in principle be generalised to account for these, and indeed other, inhibition mechanisms.

To couple the aggregation kinetics with inhibitors and liquid-liquid phase separation, we consider an approximate case where the partitioning of the aggregating monomers and the drug is close to equilibrium at all times during aggregation kinetics. This approximation is valid if these species diffuse fast compared to the characteristic timescales of aggregation and inhibitor binding (SI Sec. S1.4). Typical values of amyloid aggregation rates and condensate sizes suggest that our approximation is consistent with typical experimental conditions (SI Table S1). For simplicity, we restrict ourselves to monomers and drug molecules diffusing through the condensate interface and neglect the diffusive exchange of fibrils (Fig. 1(a)). Diffusion of fibrils is considered to be slow compared to monomers due to their large molecular weigth, which limits their diffusion via slow reptation in an environment of entangled fibrils. Under these assumptions, deviations from the equilibrium partitioning of monomers and drug molecules, caused by slow aggregation and its inhibition, are quasi-statically balanced by rapid diffusive fluxes across the interface of the liquid condensate. These fluxes ensure that the partitioning of the respective species stays approximately close to the equilibrium value at all times. Moreover, as suggested by recent inhibition experiments *in vitro* [10], the binding rate of the drug to its target is often much faster than aggregation, such that the drug binding kinetics can be considered to be quasi-statically equilibrated at all times during the aggregation kinetics (SI Sec. S2, Table S1). In this case we can capture the impact of the drug on aggregation by means of effective rate parameters (see (1f)-(1g)). By combining the above ingredients, a coupled set of kinetic equations describing inhibited aggregation in both phases *α* = I,II can be written as (SI Sec. S2):

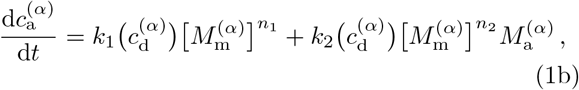

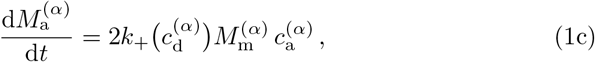

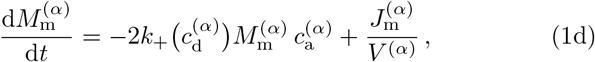

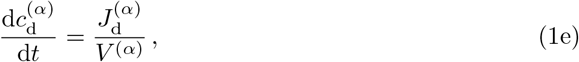

where 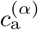 and 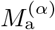 denote, respectively, the number and mass concentrations of aggregates, 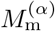 is the monomer concentration and 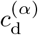 is the drug concentration in phase *α*. (1b) describes the formation of new aggregates by primary and secondary nucleation with *k*_1_, *k*_2_ being the rate constants of primary and secondary nucleation and *n*_1_, *n*_2_ denoting the respective reaction orders. Depending on the value of *n*_2_, the secondary nucleation rate describes different processes [55]: fragmentation (*n*_2_ = 0), branching (*n*_2_ = 1), and surface-catalysed secondary nucleation (*n*_2_ *>* 1) (see SI Fig. S1 for examples of amyloid forming systems and associated reaction orders). (1c) captures the buildup of aggregate mass by monomer pickup, where *k*_+_ is the rate constant for fibril elongation and the factor 2 accounts for two growing ends per aggregate. (1d) accounts for monomer depletion by fibril growth and the inter-phase flux maintaining the monomer equilibrium partitioning. (1e) describes the partitioning dynamics of the drug. Moreover, in the limit of fast inhibitor binding, the drug is in pre-equilibrium with its target and the inhibitor concentration. In this case, the on/off-binding rate constants of the drug to monomers (*i* = m), fibril ends (*i* = e) and fibril surface sites (*i* = s) enter (1) as equilibrium constants *K_i_* (SI Sec. S1.2, [31]) and the the effective aggregation rates can be written as (SI, Sec. S1.3)

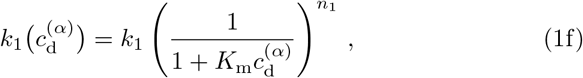

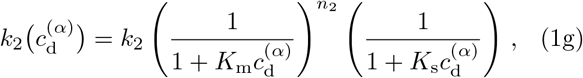

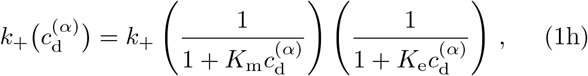

where *k*_1_ *k*_2_, *k*_+_ denote the aggregation rate constants for primary nucleation, secondary nucleation and growth in the absence of the inhibitor. Overall, our model for fibril formation in the presence of condensates and drugs has three aggregation parameters, reaction orders for primary and secondary nucleation, binding equilibrium constants for each inhibition mechanism, and three phase separation parameters: compartment volume *V*^I^, and the partitioning coefficients for drug and monomers, Γ_d_ and Γ_m_. In the following we will choose the aggregation parameters consistent with amyloid fibril formation *in vitro* [52] and vary the phase separation parameters in ranges consistent with protein condensates in *in vitro* and *in vivo*.

## RESULTS

### Condensates enhance inhibition of secondary nucleation

To study the impact of a liquid condensate on aggregation inhibition, we numerically studied equations (1) and determined asymptotic analytical solutions for the number and mass concentrations of fibrils inside and outside the liquid condensate (Fig. 2(a) and SI Sec. S2.5). These analytical solutions are valid if the timescales between the diffusion of monomers and drugs is much shorter than the comparatively slower aggregation reactions, an assumption which is realised in practice as discussed in SI Sec. 1.4. In Fig. 2(a) we show representative aggregation time courses for the number concentration of fibrils in the cleation, and with and without condensates, respectively. We consider the uninhibited, homogeneous system (no condensates, no drug) as a reference, and compare the terminal (*t*) number concentration of aggregates formed for three systems: (i) no condensates with drug, (ii) condensates, no drug and (iii) condensates with drug (Fig. 2(b)); systems (i) and (iii) have the same total drug concentration. Strikingly, while in all three systems the overall concentration of fibrils is reduced relative to the reference, we see that the system with condensates and drugs is most effective in inhibiting aggregation. Moreover, adding the drug to a heterogeneous system results in a significantly more pronounced reduction of fibrils compared to adding the same type and amount of drug to the homogeneous system (Fig. 2(b)). Therefore, the presence of liquid condensates can enhance the inhibitory action of the drug compared to the homogenous system.

**FIG. 2.**
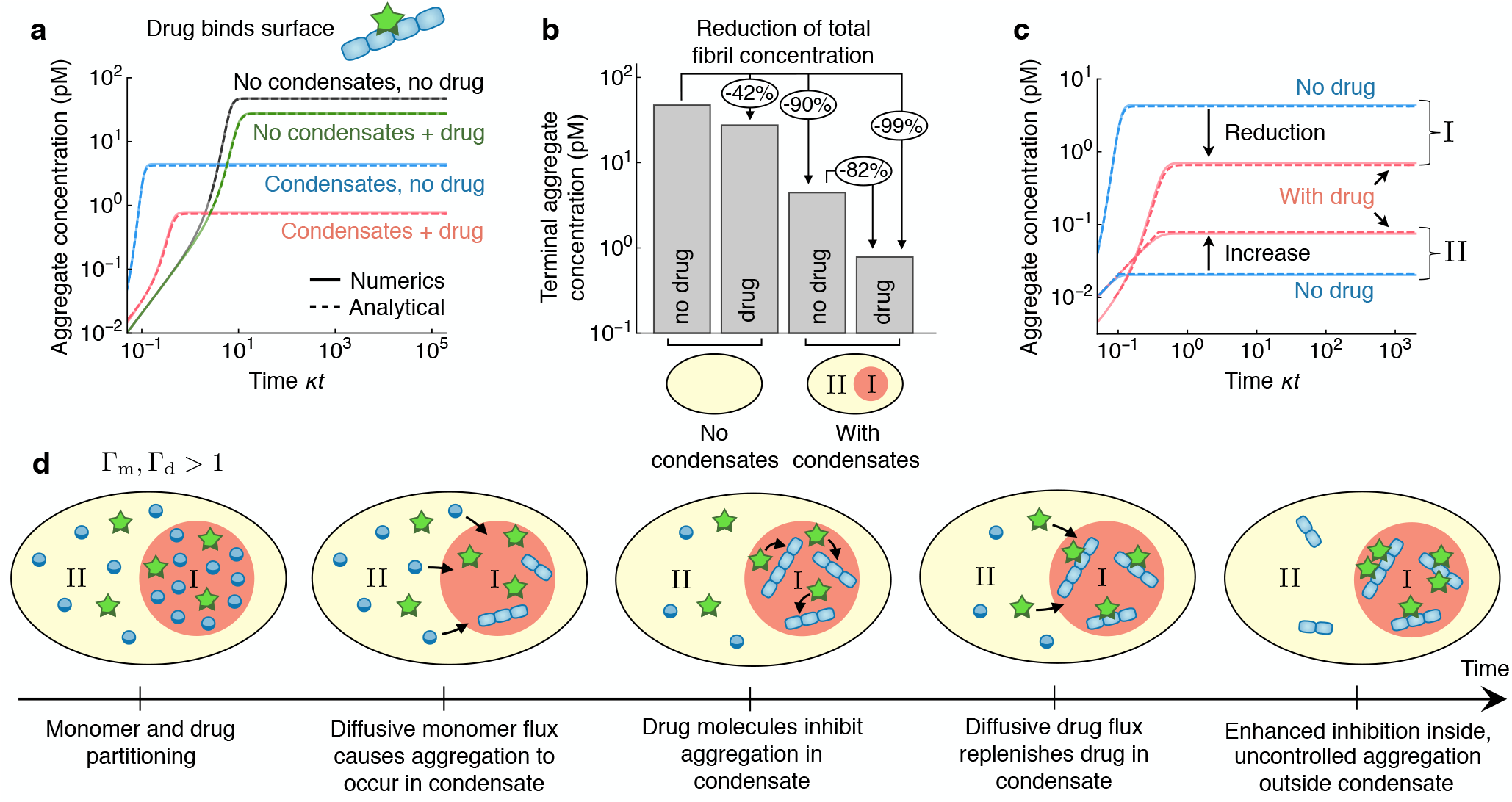
Mechanism by which liquid condensates amplify inhibition of secondary nucleation during amyloid aggregation. **(a)** Representative reaction time courses of amyloid aggregation for different systems (from top to bottom in terms of final plateau): (i) a homogeneous system without drug, (ii) a homogeneous system with a drug that inhibits secondary nucleation, and a heterogeneous system with a phase separated liquid condensate in the absence (iii) and presence (iv) of the drug. The solid lines show the numerical result from the kinetic equations (1), while the dashed lines show the analytical solution derived in this study (see SI Sec. S2.5). The plots are generated for *V*^I^*/V* = 10^*−*3^, Γ_m_ = Γ_d_ = 20 and using typical rate parameters determined previously for the A*β*42 peptide [30, 52]: 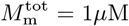, 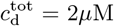, *k*_1_ = 10^*−*4^ M^*−*1^s^*−*1^, *k*_2_ = 4 *×* 10^4^ M^*−*2^s^*−*1^, *k*_+_ = 3 10^6^ M^*−*1^s^*−*1^, *n*_1_ = *n*_2_ = 2, *k*_on_ = 10^5^ M^*−*1^s^*−*1^, *k*_off_ = 10^*−*1^ s^*−*1^, *k_i_* = 10^2^ s^*−*1^. **(b)** Concentration of aggregates at the end of the aggregation reaction for the four systems (i-iv) considered in panel (a). We see that combining the drug with liquid-liquid phase separation leads to a significant decrease of the final fibril concentration compared to adding the drug to a homogeneous system. **(c)**Time course of aggregation inside (I, top part of plot) and outside (II, bottom part of plot) the liquid condensate without and with the drug. We see that adding the drug causes a reduction of aggregate concentration inside the condensate (I) but leads to an increase in aggregate concentration outside the condensate (II). **(d)**Schematic representation of the positive feedback mechanism causing enhanced inhibition inside the liquid condensate. For Γm *>* 1 aggregation occurs primarily in phase I due to the diffusive flux of monomers maintaining equilibrium. As drug molecules in phase I bind their target (Γ_d_ *>* 1), their concentration decreases. This leads to a concentration unbalance across the interface of the condensate which causes a diffusive flux of drug molecules from II to I to restore phase equilibrium. As a result, inhibition is enhanced inside the condensate (I), but the reaction proceeds in an uncontrolled fashion outside (II).

### A positive feedback mechanism underlies enhanced inhibition in the presence of condensates

To understand the mechanism underlying this enhancement effect, we consider the impact of the drug on the aggregate concentration inside (I) and outside (II) the condensate (Fig. 2(c)). We find that, while the drug inhibits aggregation in I, aggregation in II yields more aggregates in the presence of the drug compared to the case without drug. This seemingly counterintuitive effect is a direct consequence of the rapid tendency of monomers and drug components to reestablish partitioning equilibrium (Fig. 2(d)). Let us consider the case where monomers and drugs partition preferentially into the condensate (Γ_m_ *>* 1, Γ_d_ *>* 1). An enhanced concentration of monomers inside the condensate accelerates aggregation in phase I compared to the continuous phase II [38]. As aggregation occurs in phase I, further monomers diffuse from phase II to I to maintain the monomer partitioning equilibrium, causing aggregation to occur primarily in phase I [38]. Partitioning of inhibitors into the condensate then leads to an increase of binding events of drug molecules to the aggregates inside the condensate. This causes further drug molecules to diffuse from the outside into the condensate to maintain the chemical potential of inhibitors constant across the interface of the condensate. This positive feedback in phase I couples to negative feedback in phase II, where drug molecules are continuously depleted in favour of the condensate. This causes effective suppression of aggregation in I, but leaves aggregation in II potentially uncontrolled. As a result, inhibition is enhanced inside the phase (I), while it is lowered outside (II).

This coupling effect is reflected in the corresponding analytic solution to (1) which provides further mechanistic insights. In the SI Secs. S2 and S3 we show that in a system with condensates but no drug 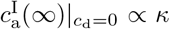 and 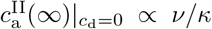, where 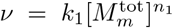 is the rate of primary nucleation and 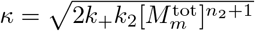 is the the effective rate of aggregate multiplication through the autocatalytic fibril cycle driven by secondary nucleation and growth [52] (Fig. 1(b)). We see that the key mechanism relies on the coupling of the autocatalytic fibril cycles inside and outside through the diffusive monomer flux (Fig. 1(c-d)). For Γ_m_ *>* 1, faster autocatalytic proliferation of aggregates in phase I promotes diffusion of monomers from II to I; the autocatalytic fibril cycle in phase I thus acts as a sink of monomers in phase II. Indeed, increasing *κ* rises 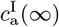 but lowers 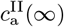. Thus, the faster the autocatalytic fibril cycle in I, the greater is the enrichment of aggregates in the condensate compared to II. Partitioning the drug in the condensate (Γ_d_ *>* 1) suppresses fibril autocatalysis in I, hence reducing the monomer flux from II to I and decreasing aggregate enrichment in I. In particular, we find (derivation see SI Sec. S3):

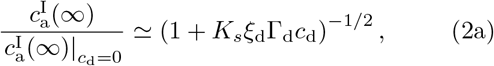

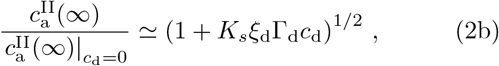

where the impact of condensate volume is captured by the partitioning degree *ξ*_d_ = 1*/*[1 + (Γ_d_ - 1)*V*^I^*/V*] (in terms of *ξ*_d_ the drug concentration in phase I is given as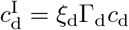. We see from (2) that upconcentrating drug molecules in phase I by increasing Γ_d_ causes effective inhibition inside phase I, while in phase II, the aggregate concentration is increased compared to the uninhibited kinetics. To understand which of these competing effects dominate we need to quantify how the phase separation parameters of volume and partitioning coefficients actually affect the total amount of aggregates in the system.

### Physical conditions for enhanced inhibition of secondary nucleation by condensates

We introduce an enhancement function 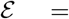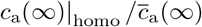 to compare the final aggregate concentration formed in a homogeneous system in the presence of the drug, *c*_a_(*∞*)*|*_homo_, with the average aggregate concentration formed with the same amount of drug in the phase-separated system, 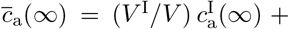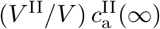 (SI Sec. S3). For *E >* 1 the condensate enhances inhibition in the entire system compared to a homogeneous solution, since the system with drop and drugs yields fewer aggregates than the system without droplets but with drugs. For large condensates (*V*^I^ ≃ *V*), we find that inhibition is enhanced for Γ_d_ ≫ 1. For Γ_d_ ≪ 1, we find ε *<* 1, i.e. the phase-separated system yields more aggregates than the homogeneous one. Enhanced inhibition therefore occurs if both the inhibitor and the target are concentrated inside the condensate (colocalization). As *V*^I^ is lowered, the contribution of 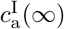 to the average concentration 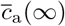 is reduced, making the system more susceptible to uncontrolled aggregation in II. Thus, in the limit of small condensates the drug should preferably partition outside the condensate (Γ_d_ ≪ 1). Overall, our results suggest that that there are two competing effects at play: effective enhancement of inhibition by the condensate requires colocalization of the drug with the aggregates inside the condensate while avoiding uncontrolled aggregation outside. Consistently, the drug should partition in phase I for large condensates, while for smaller droplets the drug should suppress aggregation outside the condensate.

### Optimal enhancement

This qualitative change of mechanism with condensate volume raises the question of which system (with/without drugs and with/without droplets; Fig. 2(b)) yields the most effective reduction of aggregate concentration. To this end, we compare the terminal fibril concentrations as a function of drug partitioning Γ_d_ for three different condensate volumes *V* ^I^*/V* and for the different systems (Fig. 3(b)): (i) no droplets with drug, (ii) droplets, no drug and (iii) droplets with drug. For large condensates the optimal system depends on Γ_d_: If Γ_d_ ≪ 1, the optimal choice is to add the drug to a homogeneous solution, while if the drug is enriched in the condensate, Γ_d_ ≫ 1, the most effective suppression of aggregation occurs in the phase-separated system in the presence of the drug. For intermediate condensate volumes, adding the drug to the heterogeneous system is the best strategy for all values of Γ_d_. By contrast, in the limit of smaller condensates adding the drug to the phase-separated system causes an *increase* of the overall fibril concentration due to uncontrolled aggregation outside the drop in phase II. In this case, the optimal solution is the phase-separated system in the absence of the drug. The resulting phase diagram, shown in Fig. 3(c), thus provides a practical strategy for choosing the most effective system to suppress aggregation depending on the phase-separation parameters.

**FIG. 3.**
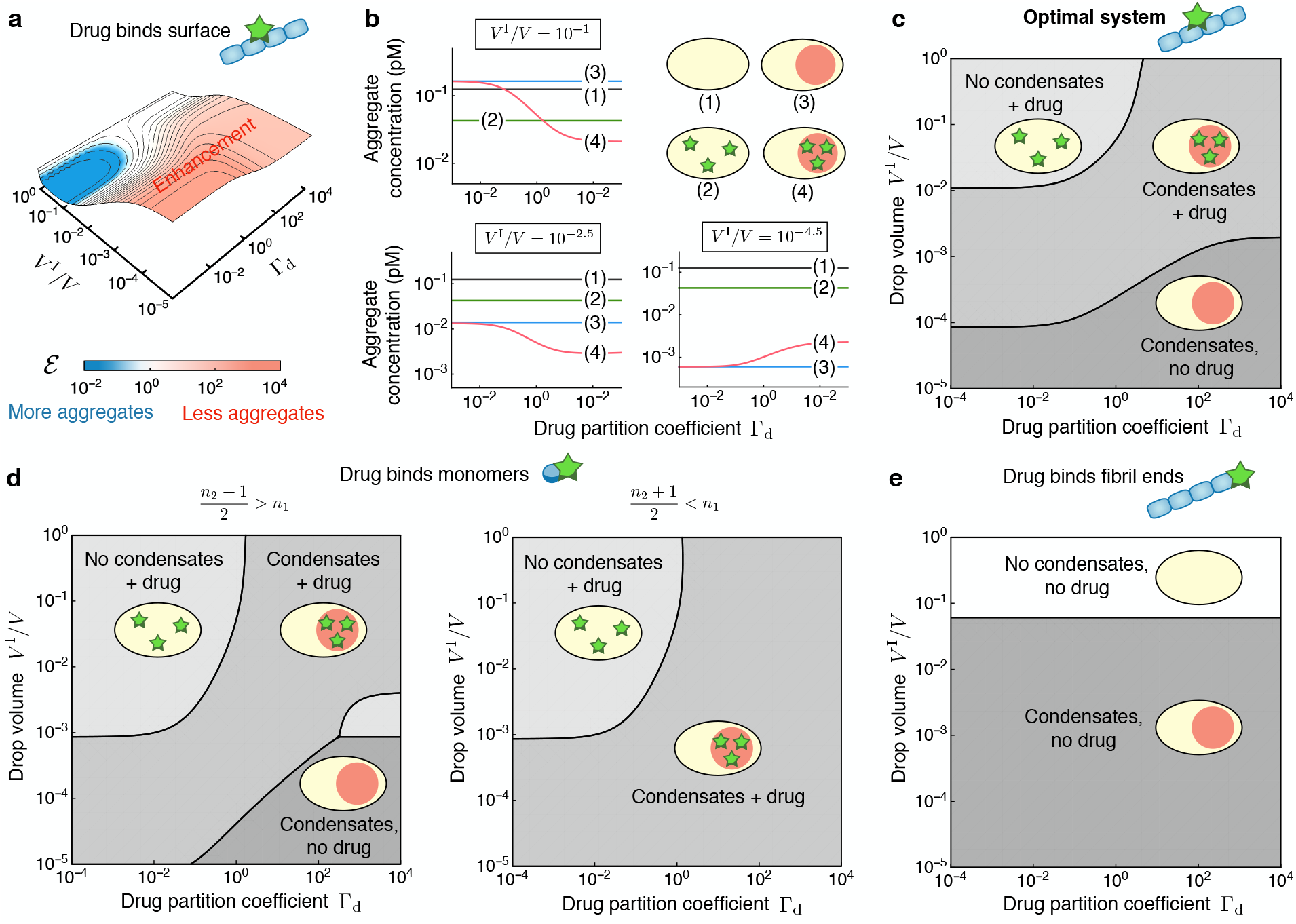
Conditions for enhanced aggregation inhibition mediated by liquid condensates and phase diagram of optimal system for inhibiting aggregation. **(a)** Enhancement function 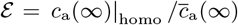 with respect to drop volume, *V*^I^*/V*, and equilibrium drug partition coefficient, Γ_d_ (for fixed Γ_m_ and drug concentration). We separate regions where liquid droplets enhance (ε*>* 1, red) or reduce (ε*<* 1, blue) the inhibitory action of a drug that binds fibril surface sites. The contour lines of the enhancement function ε are shown in logarithmic scale in steps of 10^0.5^. **(b)** Comparison of terminal aggregate concentrations formed for different systems ((1) no droplets, (2) no droplets + drug, (3) droplets, (4) droplets + drug) against drug partitioning Γ_d_. The plots are shown for fixed drop volume: *V*^I^*/V* = 10^*−*1^ (top), *V*^I^*/V* = 10^*−*2.5^ (middle) and *V*^I^*/V* = 10^*−*4.5^ (bottom). **(c)**Phase diagrams indicating the system that yields the most effective reduction of the total aggregate concentration depending on drop volume *V*^I^*/V* and drug partitioning Γ_d_ for an inhibitor of secondary nucleation. The parameters are the same as in Fig. 2 except for Γ_m_ = 10, 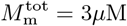 and 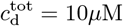. **(d-e)**Phase diagrams indicating the system that yields the most effective reduction of the total aggregate concentration for an inhibitor that binds monomers **(d)**or fibril ends **(e)**.

### Role of inhibition mechanism

We next sought to understand how the results for an inhibitor of secondary nucleation differ from other inhibition mechanisms. For example, for an inhibitor that binds aggregating monomers, we find (derivation see SI Sec. S3):

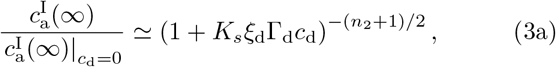

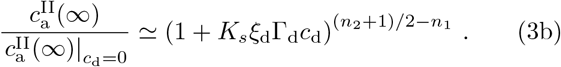

We see that the drug has opposite effects on the aggregate concentration in II depending on the sign of the exponent (*n*_2_ + 1)*/*2 − *n*_1_. Here, (*n*_2_ + 1)*/*2 is the effective reaction order of the autocatalytic fibril cycle[**?**] and *n*_1_ is the reaction order of primary nucleation. Thus, when *n*_1_ *<* (*n*_2_ + 1)*/*2 the drug inhibits the autocatalytic fibril cycle in I much more effectively than primary nucleation in II (Fig. 3(d)). This reduces the aggregate concentration in I but causes uncontrolled aggregation in II, similar to what we found for an inhibitor of secondary nucleation. By contrast, when *n*_1_ *>* (*n*_2_ + 1)*/*2 the drug inhibits primary nucleation in II much more effectively than the autocatalytic proliferation of fibrils in I. This causes effective suppression of aggregation in both phases, because aggregation in II never gets out of control. Crucially this observation implies that when the drug is an effective inhibitor of primary nucleation, adding the drug to the phase-separated system is always the optimal strategy (Fig. 3(d)). However, for drugs inhibiting growth at fibril ends, the system with droplets but without drug is always the optimal one (Fig. 3(e)). This effect is because binding the fibril ends increases the fibril number concentration.

### Condensates increase potency of aggregation inhibitors

Our theory predicts under which conditions condensates enhance aggregation inhibition compared to a homogeneous system. This indicates that an equivalent degree of inhibition compared to the homogeneous system can be realised using a smaller amount of drug in the presence of condensates. In other words liquid condensates can enhance the potency of a drug. To illustrate this effect, we define for both, the homogeneous and the phase-separated system, a drug-response curve 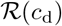 given as the relative difference between asymptotic fibril concentrations with and without drug, 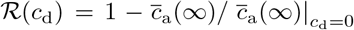 (Fig. 4a). On the basis of the response function, we introduce the drug potency 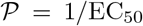 as the inverse of the drug concentration necessary to suppress 50% of the aggregates, denoted as EC_50_. The smaller EC_50_, the more potent is the drug. In Fig. 4b, we show the potency for two inhibition strategies that exhibit distinct phase diagrams for the optimal system to inhibit aggregation (Fig. 3). For an inhibitor of secondary nucleation we see that the potency increases with decreasing drop volume in the limit of large condensates. For small condensates, the optimal system becomes the phase-separated system without drug (Fig. 3c); correspondingly, the potency decreases due to uncontrolled aggregation in phase II. For an inhibitor that binds monomers with *n*_1_ *<* (*n*_2_ + 1)*/*2, the behavior is fundamentally different. The potency increases with decreasing drop volume reaching a plateau. This effect is in line with the prediction that the phase-separated system with drug is the optimal one for all values of *V*^I^*/V* (Fig. 3d). We then calculate the ratio between EC_50_ values in the absence and in the presence of condensates, respectively, which we refer to as relative potency (SI Sec. S4):

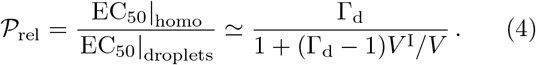

**FIG. 4.**
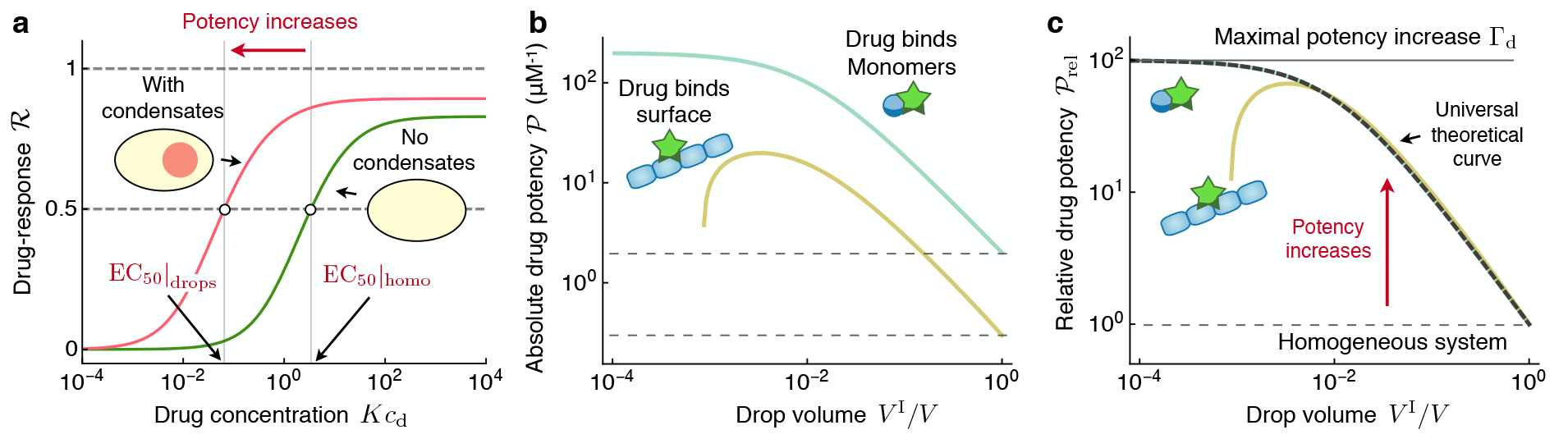
Liquid condensates increase the potency of protein aggregation inhibitors. **(a)** Drug-response curve 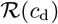 in the absence and in the presence of a liquid condensate. The presence of the liquid condensate increases the potency by reducing EC_50_. The plot is shown for an inhibitor of secondary nucleation using the same parameters as in Fig. 3c except for Γ_d_ = 10^2^ and *V*^I^*/V* = 10^*−*2^. **(b)** Potency of inhibitors that bind fibril ends (orange) or monomers (with *n*_1_ = 1 and *n*_2_ = 2, blue) against drop volume for Γ_d_ = 10^2^. The potency is defined as 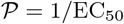 of the respective system. The dashed lines indicate the potency of the respective homogeneous system. **(c)**Relative drug potency, defined as 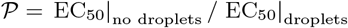, collapses onto a universal curve, (4), at least for large condensates (thick dashed line). For small condensates, the potency of the inhibitor of secondary nucleation decreases due to uncontrolled aggregation in phase II.

Strikingly, we find that while the absolute value of the potency differs among the different inhibition strategies, the relative potency is independent of the specific mechanism of inhibition, at least under the conditions when the phase-separated system with drug is the optimal system (Fig. 4c); 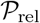 thus represents a universal relationship for how drugs affect aggregation in the presence of condensates. The relative potency 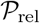 increases from unity for large condensates to a value equal to the drug partitioning Γ_d_ when the condensate volume is decreased (Fig. 4c). Hence, reducing the volume of the condensate *V* ^I^*/V* and/or increasing the drug enrichment factor Γ_d_ lead to a strong increase of the relative potency. This increase corresponds to a strong reduction of the amount of drug necessary to inhibit aggregation.

## DISCUSSION AND CONCLUSIONS

Heterogeneous environments have much potential in regulating biochemical reactions [32–35]. Our present study shows that liquid-like condensates can strongly affect drug-mediated inhibition of amyloid aggregation by providing such a heterogeneous environment composed of two coexisting phases. A key finding is that the potency of amyloid fibril inhibitors (chaperones, synthetic drugs) can be increased due to the presence of a liquid condensate. This increase in potency implies that a significantly smaller drug dose is required to equally inhibit aggregation compared to the corresponding homogeneous system. We find that this potency increase follows a universal trend with partitioning coefficient and condensate volume which is independent of the specific inhibition mechanism. Instead the potency depends on the generic colocalization of the inhibitor with its target within the liquid condensate leading to a positive feedback on the inhibition of aggregates.

To experimentally scrutinize our key findings we propose considering protein-rich condensates which recruit aggregation-prone monomers as well as aggregation inhibitors (chaperones or synthetic drugs) and determining the response as a function of drug/chaperone concentration with and without condensates (Fig. 4a).

In living cells, the colocalization of drug/chaperones and aggregates may reduce the toxic effects on the organism by preventing drug/chaperone molecules from interacting with other sensitive cellular domains. Thus, our finding of an enhanced drug potency due to the presence of condensates suggests revisiting nominally efficacious drugs that have been disregarded in a homogeneous setting due to their high toxicity and shift the focus to how drugs act specifically in the context of a spatially organized, intra-cellular environment.

## Acknowledgments

We thank Sudarshana Laha for valuable feedback on the Supplementary Material. We also thank Simon Alberti, Titus Franzmann, and Paolo Arosio, for very helpful comments on the manuscript. We acknowledge support from the Swiss National Science foundation (TCTM), Peterhouse, Cambridge (TCTM). We also thank the visitor programme of the Max Plack Institute for the Physics of Complex system for hosting TCTM and providing a stimulating research environment.

## Notes

### Competing Interest Statement

The authors have declared no competing interest.

