## Supplementary Information for "Liquid condensates increase potency of amyloid fibril inhibitors"

**S3 Calculation of enhancement function $\mathcal{E}$** **11**

**S4 Potency increase** **12**

**S5 Glossary** **13**

### S1. Kinetic equations for inhibition of amyloid aggregation in the presence of liquid drops

**S1.1. Homogeneous system without aggregation-inhibiting drug.** In the absence of a liquid compartment and of an inhibitor, the kinetics of protein aggregation into amyloid fibrils can be captured by a set of coupled differential equations for three coarse-grained fields, which correspond to commonly accessible experimental observables (1–4):

$$
\begin{aligned}
c_\mathrm{a}(t) &= \text{aggregate number concentration}\,,\\
M_\mathrm{a}(t) &= \text{aggregate mass concentration}\,,\\
M_\mathrm{m}(t) &= \text{monomer concentration}\,.
\end{aligned}
$$

In the absence of a drug that affects protein aggregation, the fundamental equations in terms of these three coarse-grained fields read (1–4):

$$\frac{\mathrm{d}c_\mathrm{a}(t)}{\mathrm{d}t} = k_1 M_\mathrm{m}(t)^{n_1} + k_2 M_\mathrm{m}(t)^{n_2} M_\mathrm{a}(t)\,, \qquad \text{[S1a]}$$

$$\frac{\mathrm{d}M_\mathrm{a}(t)}{\mathrm{d}t} = 2k_+ M_\mathrm{m}(t) c_\mathrm{a}(t) = -\frac{\mathrm{d}M_\mathrm{m}(t)}{\mathrm{d}t}\,, \qquad \text{[S1b]}$$

where parameters are

$$
\begin{aligned}
k_1 &= \text{rate constant for primary nucleation}\,,\\
k_2 &= \text{rate constant for secondary nucleation}\,,\\
k_+ &= \text{rate constant for aggregate elongation (growth)}\,,\\
n_1 &= \text{reaction order for primary nucleation}\,,\\
n_2 &= \text{reaction order for secondary nucleation}\,.
\end{aligned}
$$

| **a** Amyloid system | $n_1$ | $n_2$ | Secondary nucleation mechanism | Ref |
|---|---|---|---|---|
| Ure2p | 2 | 0 | Fragmentation | [5] |
| Actin + Arp2/3 | 3 | 1 | Branching | [5] |
| Aβ40, Aβ42 | 2 | 2 | Surface-catalysed | [8] |
| Amylin | 8 | 4 | Surface-catalysed | [5] |

| **b** Inhibitor | Amyloid system | Target species | Inhibitor type | Ref |
|---|---|---|---|---|
| Hsp70 Ss1 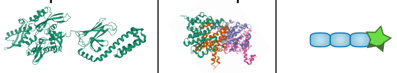 | Ure2p | Fibril ends | Molecular chaperone | [10] |
| proSP-C Brichos 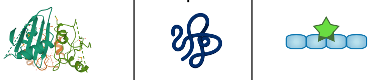 | Aβ42 | Fibril surface | Molecular chaperone | [10] |
| αB-crystallin 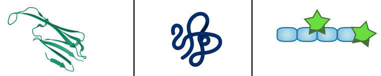 | Aβ42 | Fibrils | Molecular chaperone | [10] |
| 10074-G5 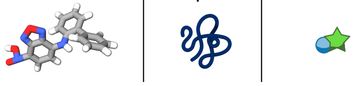 | Aβ42 | Monomers | Small molecule | [13] |

**Fig. S1. Examples of amyloid forming protein systems and inhibitors. (a)** Representative examples of different amyloid forming systems and associated reaction orders for primary and secondary nucleation $(n_1, n_2)$. **(b)** Examples of inhibitors of amyloid formation and associated target species.

See Fig. S1(a) for representative examples of amyloid forming protein systems and for associated reaction orders of primary and secondary nucleation. Note that the total concentration of monomers $M_{\mathrm{m}}^{\mathrm{tot}}$ (free and aggregated) is conserved at all times

$$M_{\mathrm{m}}^{\mathrm{tot}} = M_{\mathrm{m}}(t) + M_{\mathrm{a}}(t). \qquad \text{[S2]}$$

***Aggregation time course.*** As discussed in detail in Ref. (4), the solution to Eq. (S1) exhibits a sigmoidal shape in time. When secondary nucleation dominates over primary nucleation, the aggregate number and mass concentrations initially grow exponentially with time with rate

$$\kappa_0 = \sqrt{2k_+k_2[M_{\mathrm{m}}^{\mathrm{tot}}]^{n_2+1}}.$$

$\kappa_0$ is thus a combined rate of aggregate proliferation through secondary nucleation and elongation which sets the basic unit of time of the aggregation reaction. After the initial exponential growth phase, the kinetic profile saturates due to depletion of monomers. An analytical solution to the kinetics has been obtained previously (5):

$$\frac{M_{\mathrm{a}}(t)}{M_{\mathrm{m}}^{\mathrm{tot}}} = 1 - \frac{M_{\mathrm{m}}(t)}{M_{\mathrm{m}}^{\mathrm{tot}}} = 1 - \left[1 + \frac{\lambda_0^2}{2\kappa_0^2}e^{\kappa_0 t}\right]^{-\theta}, \qquad \text{[S3a]}$$

$$\frac{c_{\mathrm{a}}(t)}{c_{\mathrm{a}}(\infty)} = \left[1 + \frac{2\kappa_0^2}{\lambda_0^2}e^{-\kappa_0 t}\right]^{-1}, \qquad \text{[S3b]}$$

where the asymptotic number concentration of aggregates is

$$c_{\mathrm{a}}(\infty) = \frac{1}{2k_+}\sqrt{\kappa_0^2\theta^2 + \frac{2\lambda_0^2}{n_1}} = \frac{\kappa_0\theta}{2k_+}\sqrt{1 + \frac{2w}{n_1\theta}}, \qquad \text{[S3c]}$$

and $\theta = \sqrt{2/[n_2(n_2+1)]}$, $\lambda_0 = \sqrt{2k_+k_1[M_{\mathrm{m}}^{\mathrm{tot}}]^{n_1}}$. The parameter $w$

$$w = \frac{k_1\,(M_{\mathrm{m}}^{\mathrm{tot}})^{n_1-n_2-1}}{k_2\theta} \qquad \text{[S3d]}$$

measures the ratio between the rates of primary and secondary nucleation. For aggregating systems where secondary nucleation pathways dominate over the primary nucleation process for the formation of new filaments, we have $w \ll 1$. Typical values for $w$ are in fact $w \simeq 10^{-2}$ for the yeast prion Ure2p (6), $w \simeq 10^{-3}$ for sickle-cell haemoglobin (Hsb) (7), $w \simeq 10^{-5}$ for the amyloid-$\beta$ (A$\beta$) peptide (8), and $w \simeq 10^{-7}$ for the islet amyloid polypeptide (IAPP) (9). We can thus focus on the limit $w \ll 1$, such that Eq. (S3c) becomes:

$$c_\mathrm{a}(\infty) \simeq \frac{\kappa_0 \theta}{2k_+}. \tag{S3e}$$

**S1.2. Homogeneous system with aggregation-inhibiting drug.** The presence of an inhibitor can affect different microscopic events of aggregation in different ways (see Fig. S1(b) for representative examples of inhibitors and targeted species). First, the inhibitor can bind to monomers and inhibit nucleation and growth. Second, it can bind to fibril ends and inhibits elongation (growth). And third, the inhibitor can bind to fibril surfaces and inhibit secondary nucleation. To describe these binding processes additional coarse-grained fields are required which account for the free and bound components, respectively:

$$\begin{aligned}
c_\mathrm{a,f}(t) &= \text{free aggregate number concentration}\,,\\
c_\mathrm{a,b}(t) &= \text{bound aggregate number concentration}\,,\\
M_\mathrm{a,f}(t) &= \text{free aggregate mass concentration}\,,\\
M_\mathrm{a,b}(t) &= \text{bound aggregate mass concentration}\,,\\
M_\mathrm{m,f}(t) &= \text{free monomer concentration}\,,\\
M_\mathrm{m,b}(t) &= \text{bound monomer concentration}\,,\\
c_\mathrm{d}(t) &= \text{(free) inhibitor concentration}\,.
\end{aligned}$$

The fundamental equations that describe the kinetics of protein aggregation in the presence of an inhibitor are (10, 11):

$$\frac{\mathrm{d}c_\mathrm{a,f}(t)}{\mathrm{d}t} = k_1 M_\mathrm{m,f}(t)^{n_1} + k_2 M_\mathrm{m,f}(t)^{n_2} M_\mathrm{a,f}(t) - k_\mathrm{on,e} c_\mathrm{a,f}(t) c_\mathrm{d}(t) + k_\mathrm{off,e} c_\mathrm{a,b}(t)\,, \tag{S4a}$$

$$\frac{\mathrm{d}M_\mathrm{a,f}(t)}{\mathrm{d}t} = 2k_+ M_\mathrm{m,f}(t) c_\mathrm{a,f}(t) - k_\mathrm{on,s} M_\mathrm{a,f}(t) c_\mathrm{d}(t) + k_\mathrm{off,s} M_\mathrm{a,b}(t)\,, \tag{S4b}$$

$$\frac{\mathrm{d}M_\mathrm{m,f}(t)}{\mathrm{d}t} = -2k_+ M_\mathrm{m,f}(t) c_\mathrm{a,f}(t) - k_\mathrm{on,m} M_\mathrm{m,f}(t) c_\mathrm{d}(t) + k_\mathrm{off,m} M_\mathrm{m,b}(t)\,, \tag{S4c}$$

for free species,

$$\frac{\mathrm{d}c_\mathrm{a,b}(t)}{\mathrm{d}t} = k_\mathrm{on,e} c_\mathrm{a,f}(t) c_\mathrm{d}(t) - k_\mathrm{off,e} c_\mathrm{a,b}(t)\,, \tag{S4d}$$

$$\frac{\mathrm{d}M_\mathrm{a,b}(t)}{\mathrm{d}t} = k_\mathrm{on,s} M_\mathrm{a,f}(t) c_\mathrm{d}(t) - k_\mathrm{off,s} M_\mathrm{a,b}(t)\,, \tag{S4e}$$

$$\frac{\mathrm{d}M_\mathrm{m,b}(t)}{\mathrm{d}t} = k_\mathrm{on,m} M_\mathrm{m,f}(t) c_\mathrm{d}(t) - k_\mathrm{off,m} M_\mathrm{m,b}(t)\,, \tag{S4f}$$

for bound species and

$$\frac{\mathrm{d}c_\mathrm{d}(t)}{\mathrm{d}t} = -\frac{\mathrm{d}c_\mathrm{a,b}(t)}{\mathrm{d}t} - \frac{\mathrm{d}M_\mathrm{a,b}(t)}{\mathrm{d}t} - \frac{\mathrm{d}M_\mathrm{m,b}(t)}{\mathrm{d}t}\,, \tag{S4g}$$

for the drug concentration. Here, the binding rates are

$$\begin{aligned}
k_\mathrm{on,e} &= \text{on-rate constant for inhibitor binding to aggregate ends}\,,\\
k_\mathrm{off,e} &= \text{off-rate constant for inhibitor binding to aggregate ends}\,,\\
k_\mathrm{on,s} &= \text{on-rate constant for inhibitor binding to aggregate surface}\,,\\
k_\mathrm{off,s} &= \text{off-rate constant for inhibitor binding to aggregate surface}\,,\\
k_\mathrm{on,m} &= \text{on-rate constant for inhibitor binding to monomers}\,,\\
k_\mathrm{off,m} &= \text{off-rate constant for inhibitor binding to monomers}\,.
\end{aligned}$$

Eqs. (S4) couple to the conservation of total protein mass $M_{\mathrm{m}}^{\mathrm{tot}}$ which implies:

$$M_{\mathrm{m}}^{\mathrm{tot}} = M_{\mathrm{m,f}}(t) + M_{\mathrm{m,b}}(t) + M_{\mathrm{a,f}}(t) + M_{\mathrm{a,b}}(t)\,. \tag{S5}$$

For convenience, we introduce total aggregate number, total aggregate mass and total monomer concentrations as sums of the respective free and bound components:

$$c_{\mathrm{a}}(t) = c_{\mathrm{a,f}}(t) + c_{\mathrm{a,b}}(t)\,, \tag{S6a}$$
$$M_{\mathrm{a}}(t) = M_{\mathrm{a,f}}(t) + M_{\mathrm{a,b}}(t)\,, \tag{S6b}$$
$$M_{\mathrm{m}}(t) = M_{\mathrm{m,f}}(t) + M_{\mathrm{m,b}}(t)\,. \tag{S6c}$$

**S1.3. Phase-separated system.** In the presence of a liquid droplet, we have to consider different factors:

- Aggregation occurs inside each compartment ($\alpha$ = I,II) as a result of primary nucleation, secondary nucleation and growth, as described in the previous sections.
- Monomers and drug molecules partition inside (I) and outside (II) of the droplet. This gives rise to a diffusive exchange of monomers and drug molecules between the two compartments. Here we restrict ourselves to monomers and drug molecules diffusing through the compartment interface and neglect the diffusive exchange of aggregates. This is justified if diffusion of monomers and drug molecules is fast while aggregates hardly diffuse. We assume that aggregates bound to the drug hardly diffuse as well.

***S1.3.1. Monomer and drug partitioning.*** The partitioning coefficients of free monomers, bound monomers and drug molecules are defined as (12):

$$\Gamma_{\mathrm{m,f}} = \left.\frac{M_{\mathrm{m,f}}^{\mathrm{I}}}{M_{\mathrm{m,f}}^{\mathrm{II}}}\right|_{\mathrm{eq}}\,, \quad \Gamma_{\mathrm{m,b}} = \left.\frac{M_{\mathrm{m,b}}^{\mathrm{I}}}{M_{\mathrm{m,b}}^{\mathrm{II}}}\right|_{\mathrm{eq}}\,, \quad \Gamma_{\mathrm{d}} = \left.\frac{c_{\mathrm{d}}^{\mathrm{I}}}{c_{\mathrm{d}}^{\mathrm{II}}}\right|_{\mathrm{eq}}\,, \tag{S7}$$

where the subscript "eq" refers to the respective concentrations at partitioning equilibrium. The equilibrium partitioning coefficients for monomers and drug molecules can be calculated from the Flory-Huggins free energy for a mixture of free/bound monomers, inhibitor molecules and components A and B:

$$f = \frac{k_B T}{\nu}\Big[\phi_A \ln(\phi_A) + \phi_B \ln(\phi_B) + \sum_i \nu c_i \ln(\nu_i c_i) \\ + \chi_{AB}\phi_A\phi_B + \sum_i (\chi_{Ai}\phi_A \nu_i c_i + \chi_{Bi}\phi_B \nu_i c_i) + \sum_{i,j} \chi_{ij}\nu_i\nu_j c_i c_j\Big]\,, \tag{S8}$$

where $\phi_A$ and $\phi_B$ are the volume fractions of A and B and the index $i$ stands for free monomers, bound monomers and inhibitor molecules. $\nu$ is the molecular volume of A and B, $\nu_i$ is the molecular volume of $i$ = free monomers, bound monomers or inhibitor molecules. $\chi_{ij}$ is the interaction strength parameter between species $i$ and $j$. Assuming that free monomers, bound monomers and inhibitor molecules are dilute ($\sum_i \nu_i c_i \ll 1$) and using $\phi_A = 1-\phi_B-\sum_i \nu_i c_i \simeq 1-\phi_A$, we obtain after expanding $f$ at leading order in $\nu_i c_i$:

$$f \simeq \frac{k_B T}{\nu}\Big[\phi \ln(\phi) + (1-\phi)\ln(1-\phi) + \chi_{AB}\phi(1-\phi) \\ + \sum_i \nu_i c_i \Big(\frac{\nu}{\nu_i}\ln(\nu_i c_i) - \ln(1-\phi) + (\chi_{Ai} - \chi_{Bi} - \chi_{AB})\phi + \chi_{Bi} - 1\Big)\Big]\,, \tag{S9}$$

where for simplicity we set $\phi = \phi_A$. At equilibrium, we set the chemical potentials of A and B as well as species $i$ equal across the droplet boundary

$$\mu(\phi^{\mathrm{I}}, c_i^{\mathrm{I}}) = \mu(\phi^{\mathrm{II}}, c_i^{\mathrm{II}}), \tag{S10}$$
$$\mu_i(\phi^{\mathrm{I}}, c_i^{\mathrm{I}}) = \mu_i(\phi^{\mathrm{II}}, c_i^{\mathrm{II}}), \tag{S11}$$

where $\mu = \nu\,\partial f/\partial\phi$ and $\mu_i = \partial f/\partial c_i$. Eq. (S11) yields the phase equilibrium of a binary mixture, i.e. the dilute species $i$ have a negligible impact on the phase-separation of A and B . Eq. (S11), together with Eq. (S10), yields the equilibrium partitioning of species $i$ = free monomers, bound monomers or inhibitor molecules:

$$\Gamma_i = \left.\frac{c_i^{\mathrm{I}}}{c_i^{\mathrm{II}}}\right|_{\mathrm{eq}} \simeq \exp\left[\frac{\nu_i}{\nu}(\phi^{\mathrm{I}} - \phi^{\mathrm{II}})\,(\chi_{B,i} - \chi_{A,i})\right]\,, \tag{S12}$$

where $\phi^{\mathrm{I}} - \phi^{\mathrm{II}}$ is the degree of phase separation.

**S1.3.2. Inter-compartment flux of monomers and drug molecules near equilibrium.** A deviation of monomer or drug partitioning from equilibrium causes diffusive fluxes through the compartment interface. We can compute this flux of species $i$ assuming that the droplet is a sphere of radius $R$ and upon expanding the chemical potential Eq. (S11) at leading order around the equilibrium concentrations $c_i^{\mathrm{I}}|_{\mathrm{eq}}$ and $c_i^{\mathrm{II}}|_{\mathrm{eq}}$ (12):

$$J^{\mathrm{I}}(c_i) = -J^{\mathrm{II}}(c_i) = -J(c_i) = -k_i \left( c_i^{\mathrm{I}} - \Gamma(c_i) c_i^{\mathrm{I}} \right) , \quad \text{[S13]}$$

where $k_i = 4\pi R D_i$ with $D_i$ being the diffusion coefficient of species $i$. Explicitly:

$$J_{\mathrm{m,f}} = k_{\mathrm{m,f}} \left( M_{\mathrm{m,f}}^{\mathrm{I}}(t) - \Gamma_{\mathrm{m,f}} M_{\mathrm{m,f}}^{\mathrm{II}}(t) \right) , \quad \text{[S14a]}$$

$$J_{\mathrm{m,b}} = k_{\mathrm{m,b}} \left( M_{\mathrm{m,b}}^{\mathrm{I}}(t) - \Gamma_{\mathrm{m,b}} M_{\mathrm{m,b}}^{\mathrm{II}}(t) \right) , \quad \text{[S14b]}$$

$$J_{\mathrm{d}} = k_{\mathrm{d}} \left( c_{\mathrm{d}}^{\mathrm{I}}(t) - \Gamma_{\mathrm{d}} c_{\mathrm{d}}^{\mathrm{II}}(t) \right) . \quad \text{[S14c]}$$

For simplicity, we will set $\Gamma_{\mathrm{m,f}} = \Gamma_{\mathrm{m,b}} =: \Gamma_{\mathrm{m}}$ throughout.

**S1.3.3. Kinetic equations in the presence of liquid compartments.** Accounting for the combined effects of aggregation kinetics and partitioning of monomers and drugs, equations (S4) become in the presence of liquid drops:

$$\frac{\mathrm{d}c_{\mathrm{a,f}}^{(\alpha)}(t)}{\mathrm{d}t} = k_1 M_{\mathrm{m,f}}^{(\alpha)}(t)^{n_1} + k_2 M_{\mathrm{m,f}}^{(\alpha)}(t)^{n_2} M_{\mathrm{a,f}}^{(\alpha)}(t) - k_{\mathrm{on,e}} c_{\mathrm{a,f}}^{(\alpha)}(t) c_{\mathrm{d}}^{(\alpha)}(t) + k_{\mathrm{off,e}} c_{\mathrm{a,b}}^{(\alpha)}(t) , \quad \text{[S15a]}$$

$$\frac{\mathrm{d}M_{\mathrm{a,f}}^{(\alpha)}(t)}{\mathrm{d}t} = 2k_+ M_{\mathrm{m,f}}^{(\alpha)}(t) c_{\mathrm{a,f}}^{(\alpha)}(t) - k_{\mathrm{on,s}} M_{\mathrm{a,f}}^{(\alpha)}(t) c_{\mathrm{d}}^{(\alpha)}(t) + k_{\mathrm{off,s}} M_{\mathrm{a,b}}^{(\alpha)}(t) , \quad \text{[S15b]}$$

$$\frac{\mathrm{d}M_{\mathrm{m,f}}^{(\alpha)}(t)}{\mathrm{d}t} = -2k_+ M_{\mathrm{m,f}}^{(\alpha)}(t) c_{\mathrm{a,f}}^{(\alpha)}(t) - k_{\mathrm{on,m}} M_{\mathrm{m,f}}^{(\alpha)}(t) c_{\mathrm{d}}^{(\alpha)}(t) + k_{\mathrm{off,m}} M_{\mathrm{m,b}}^{(\alpha)}(t) + \frac{J_{\mathrm{m,f}}^{(\alpha)}}{V^{(\alpha)}} , \quad \text{[S15c]}$$

$$\frac{\mathrm{d}c_{\mathrm{a,b}}^{(\alpha)}(t)}{\mathrm{d}t} = k_{\mathrm{on,e}} c_{\mathrm{a,f}}^{(\alpha)}(t) c_{\mathrm{d}}^{(\alpha)}(t) - k_{\mathrm{off,e}} c_{\mathrm{a,b}}^{(\alpha)}(t) , \quad \text{[S15d]}$$

$$\frac{\mathrm{d}M_{\mathrm{a,b}}^{(\alpha)}(t)}{\mathrm{d}t} = k_{\mathrm{on,s}} M_{\mathrm{a,f}}^{(\alpha)}(t) c_{\mathrm{d}}^{(\alpha)}(t) - k_{\mathrm{off,s}} M_{\mathrm{a,b}}^{(\alpha)}(t) , \quad \text{[S15e]}$$

$$\frac{\mathrm{d}M_{\mathrm{m,b}}^{(\alpha)}(t)}{\mathrm{d}t} = k_{\mathrm{on,m}} M_{\mathrm{m,f}}^{(\alpha)}(t) c_{\mathrm{d}}^{(\alpha)}(t) - k_{\mathrm{off,m}} M_{\mathrm{m,b}}^{(\alpha)}(t) + \frac{J_{\mathrm{m,b}}^{(\alpha)}}{V^{(\alpha)}} , \quad \text{[S15f]}$$

$$\frac{\mathrm{d}c_{\mathrm{d}}^{(\alpha)}(t)}{\mathrm{d}t} = -\frac{\mathrm{d}c_{\mathrm{a,b}}^{(\alpha)}(t)}{\mathrm{d}t} - \frac{\mathrm{d}M_{\mathrm{a,b}}^{(\alpha)}(t)}{\mathrm{d}t} - \frac{\mathrm{d}M_{\mathrm{m,b}}^{(\alpha)}(t)}{\mathrm{d}t} + \frac{J_{\mathrm{d}}^{(\alpha)}}{V^{(\alpha)}} , \quad \text{[S15g]}$$

where $J_i^{\mathrm{II}} = -J_i^{\mathrm{I}} = J_i$ and the partitioning fluxes $J_i$ are given in equations (S14). This is equation (1) of the main text.

**S1.4. Model validity and separation of timescales.** Eqs. S15 consider homogeneous concentrations in each of the phases I and II. For these equations to be valid, spatial gradients musty be small. This is the case when reaction slow compared to diffusion, i.e.,

$$\sqrt{\frac{D}{k_{\mathrm{react}}}} \gg V^{1/3} , \quad \text{[S16]}$$

where $V^{-1/3}$ is the system size and $k_{\mathrm{react}}$ is the reaction rate corresponding to drug binding or aggregation, respectively. Introducing the time-scale of diffusion on the system size as $\tau_{\mathrm{diff}} = V^{2/3}/D$, the condition above is equivalent to

$$\frac{\tau_{\mathrm{react}}}{\tau_{\mathrm{diff}}} \gg 1 , \quad \text{[S17]}$$

where $\tau_{\mathrm{react}} = 1/k_{\mathrm{react}}$. For our framework to be valid, there must be a separation of time-scales between diffusion and reaction (drug binding and aggregation). In addition, we consider a further time-scale separation between drug binding and aggregation, i.e., binding is fast compared to the aggregation. Typical values from *in vitro* studies suggest the validity of this time-scale separation, see Table S1. In summary, we work with the following separation of time-scales:

$$\tau_{\mathrm{diff,m}} \simeq \tau_{\mathrm{diff,d}} \ll \tau_{\mathrm{binding}}, \tau_{\mathrm{unbinding}} \ll \tau_{\mathrm{aggregation}} . \quad \text{[S18]}$$

**Table S1.** Hierarchy of timescales in our problem using parameters from *in vitro* studies.

| Process | Typical timescale | |
|---|---|---|
| Binding kinetics of inhibitor to target | $\tau_{\text{binding/unbinding}} \simeq 1/(k_{\text{on,m}} c_{\text{d}}) \simeq 1/(k_{\text{off,m}}) \simeq 1$ min (see e.g. Ref. (13)) | intermediate |
| Aggregation kinetics | $\tau_{\text{aggregation}} \simeq 1/\kappa_0 \simeq 1$ h (see e.g. Ref. (8)) | slow |

### S2. Analytical solution to aggregation kinetics

Using the mathematical methods outlined in Refs. (5, 12), including an analogy to classical mechanics, we have obtained explicit solutions to the kinetic equations Eq. (S30) valid in the limit $\Gamma_{\text{m}} \gg 1$ and when secondary nucleation processes dominate the production of new aggregates.

Using matched asymptotics analysis (14) we can integrate out the fast degrees of freedom associated with monomer/drug partitioning and inhibitor binding kinetics. We apply this procedure in two steps, by considering partitioning kinetics first (Sec. S2.1) and then discussing the effect of fast drug-binding kinetics in Sec. S2.2

**S2.1. Fast monomer/drug partitioning.** When partitioning dynamics is very fast compared to the other processes, the solution to Eq. (S15) develops in two stages.

- There is an initial layer, where the initial values of the (free/bound) monomer and drug concentrations rapidly equilibrate, through the diffusive fluxes $J_i$, to establish partitioning equilibrium set by $\Gamma_{\text{m}}$ and $\Gamma_{\text{d}}$.

- After the fast initial phase of drug and monomer equilibration in the two compartments, we enter a slow dynamical phase (slow manifold), where the system stays close to the equilibrium partitioning to leading order. The diffusive fluxes $J_i \simeq 0$ are then close to zero at all times, imposing boundary conditions on concentrations inside and outside of the compartment. In particular, in the slow manifold the following conditions

$$M^{\text{I}}_{\text{m,f}}(t) = \Gamma_{\text{m}} M^{\text{II}}_{\text{m,f}}(t), \quad M^{\text{I}}_{\text{m,b}}(t) = \Gamma_{\text{m}} M^{\text{II}}_{\text{m,b}}(t), \quad c^{\text{I}}_{\text{d}}(t) = \Gamma_{\text{d}} c^{\text{II}}_{\text{d}}(t) \qquad \text{[S19]}$$

hold at all times. The validity of these relationships is verified numerically in Fig. S2(a). We can also express these conditions in terms of the total (free/bound) monomer and drug concentrations:

$$M_{\text{m,f}}(t) = \frac{V^{\text{I}} M^{\text{I}}_{\text{m,f}}(t) + V^{\text{II}} M^{\text{II}}_{\text{m,f}}(t)}{V}, \qquad \text{[S20a]}$$

$$M_{\text{m,b}}(t) = \frac{V^{\text{I}} M^{\text{I}}_{\text{m,b}}(t) + V^{\text{II}} M^{\text{II}}_{\text{m,b}}(t)}{V}, \qquad \text{[S20b]}$$

$$c_{\text{d}}(t) = \frac{V^{\text{I}} c^{\text{I}}_{\text{d}}(t) + V^{\text{II}} c^{\text{II}}_{\text{d}}(t)}{V}. \qquad \text{[S20c]}$$

Combining Eq. (S20) with Eq. (S19), we obtain

$$M^{\text{I}}_{\text{m,f}}(t) = \xi_{\text{m}} \Gamma_{\text{m}} M_{\text{m,f}}(t), \qquad M^{\text{II}}_{\text{m,f}}(t) = \xi_{\text{m}} M_{\text{m,f}}(t), \qquad \text{[S21a]}$$

$$M^{\text{I}}_{\text{m,b}}(t) = \xi_{\text{m}} \Gamma_{\text{m}} M_{\text{m,b}}(t), \qquad M^{\text{II}}_{\text{m,b}}(t) = \xi_{\text{m}} M_{\text{m,b}}(t), \qquad \text{[S21b]}$$

$$c^{\text{I}}_{\text{d}}(t) = \xi_{\text{d}} \Gamma_{\text{d}} c_{\text{d}}(t), \qquad c^{\text{II}}_{\text{d}}(t) = \xi_{\text{d}} c_{\text{d}}(t), \qquad \text{[S21c]}$$

where

$$\xi_{\text{m}} = \frac{1}{1 + (\Gamma_{\text{m}} - 1) V^{\text{I}}/V}, \quad \xi_{\text{d}} = \frac{1}{1 + (\Gamma_{\text{d}} - 1) V^{\text{I}}/V}. \qquad \text{[S22]}$$

In the slow manifold, we can thus set diffusive fluxes to zero in Eq. (S15) and instead impose conditions Eq. (S19) and Eq. (S21) on the variables. As a result, we can solve the following equations for the dynamics inside compartment I first:

$$\frac{\mathrm{d}c^{\mathrm{I}}_{\mathrm{a,f}}(t)}{\mathrm{d}t} = k_1 M^{\mathrm{I}}_{\mathrm{m,f}}(t)^{n_1} + k_2 M^{\mathrm{I}}_{\mathrm{m,f}}(t)^{n_2} M^{\mathrm{I}}_{\mathrm{a,f}}(t) - k_{\mathrm{on,e}} c^{\mathrm{I}}_{\mathrm{a,f}}(t) c^{\mathrm{I}}_{\mathrm{d}}(t) + k_{\mathrm{off,e}} c^{\mathrm{I}}_{\mathrm{a,b}}(t)\,, \tag{S23a}$$

$$\frac{\mathrm{d}M^{\mathrm{I}}_{\mathrm{a,f}}(t)}{\mathrm{d}t} = 2k_+ M^{(\alpha)}_{\mathrm{m,f}}(t) c^{\mathrm{I}}_{\mathrm{a,f}}(t) - k_{\mathrm{on,s}} M^{\mathrm{I}}_{\mathrm{a,f}}(t) c^{(\alpha)}_{\mathrm{d}}(t) + k_{\mathrm{off,s}} M^{\mathrm{I}}_{\mathrm{a,b}}(t)\,, \tag{S23b}$$

$$\frac{\mathrm{d}M^{\mathrm{I}}_{\mathrm{m,f}}(t)}{\mathrm{d}t} = -2k_+ M^{\mathrm{I}}_{\mathrm{m,f}}(t) c^{\mathrm{I}}_{\mathrm{a,f}}(t) - k_{\mathrm{on,m}} M^{\mathrm{I}}_{\mathrm{m,f}}(t) c^{\mathrm{I}}_{\mathrm{d}}(t) + k_{\mathrm{off,m}} M^{\mathrm{I}}_{\mathrm{m,b}}(t)\,, \tag{S23c}$$

$$\frac{\mathrm{d}c^{\mathrm{I}}_{\mathrm{a,b}}(t)}{\mathrm{d}t} = k_{\mathrm{on,e}} c^{\mathrm{I}}_{\mathrm{a,f}}(t) c^{\mathrm{I}}_{\mathrm{d}}(t) - k_{\mathrm{off,e}} c^{\mathrm{I}}_{\mathrm{a,b}}(t)\,, \tag{S23d}$$

$$\frac{\mathrm{d}M^{\mathrm{I}}_{\mathrm{a,b}}(t)}{\mathrm{d}t} = k_{\mathrm{on,s}} M^{\mathrm{I}}_{\mathrm{a,f}}(t) c^{\mathrm{I}}_{\mathrm{d}}(t) - k_{\mathrm{off,s}} M^{\mathrm{I}}_{\mathrm{a,b}}(t)\,, \tag{S23e}$$

$$\frac{\mathrm{d}M^{\mathrm{I}}_{\mathrm{m,b}}(t)}{\mathrm{d}t} = k_{\mathrm{on,m}} M^{\mathrm{I}}_{\mathrm{m,f}}(t) c^{\mathrm{I}}_{\mathrm{d}}(t) - k_{\mathrm{off,m}} M^{\mathrm{I}}_{\mathrm{m,b}}(t)\,, \tag{S23f}$$

$$\frac{\mathrm{d}c^{\mathrm{I}}_{\mathrm{d}}(t)}{\mathrm{d}t} = -\frac{\mathrm{d}c^{\mathrm{I}}_{\mathrm{a,b}}(t)}{\mathrm{d}t} - \frac{\mathrm{d}M^{\mathrm{I}}_{\mathrm{a,b}}(t)}{\mathrm{d}t} - \frac{\mathrm{d}M^{\mathrm{I}}_{\mathrm{m,b}}(t)}{\mathrm{d}t}\,, \tag{S23g}$$

subject to the initial conditions $M^{\mathrm{I}}_{\mathrm{m}}(0) = \xi_{\mathrm{m}}\Gamma_{\mathrm{m}} M^{\mathrm{tot}}_{\mathrm{m}}$ and $c^{\mathrm{I}}_{\mathrm{d}}(0) = \xi_{\mathrm{d}}\Gamma_{\mathrm{d}} c_{\mathrm{d}}$. We can then use Eq. (S19) to determine the dynamics in compartment II.

**S2.2. Fast drug binding.** After having integrated out the fast timescales associated with monomer/drug partitioning, we can focus on the timescale separation between drug binding and aggregation kinetics. In the limit when drug-binding kinetics is fast compared to aggregation, we can set

$$\frac{\mathrm{d}c^{(\alpha)}_{\mathrm{a,b}}(t)}{\mathrm{d}t} = \frac{\mathrm{d}M^{(\alpha)}_{\mathrm{a,b}}(t)}{\mathrm{d}t} = \frac{\mathrm{d}M^{(\alpha)}_{\mathrm{m,b}}(t)}{\mathrm{d}t} \simeq 0\,. \tag{S24}$$

These conditions imply

$$\frac{\mathrm{d}c^{(\alpha)}_{\mathrm{d}}(t)}{\mathrm{d}t} \simeq 0\,, \tag{S25}$$

i.e. the drug concentration is approximately constant. Then we find:

$$0 = k_{\mathrm{on,e}} c^{(\alpha)}_{\mathrm{a,f}}(t) c^{(\alpha)}_{\mathrm{d}}(t) - k_{\mathrm{off,e}} c^{(\alpha)}_{\mathrm{a,b}}(t) \quad \Rightarrow \quad c^{(\alpha)}_{\mathrm{a,b}}(t) = K_{\mathrm{e}}\, c^{(\alpha)}_{\mathrm{a,f}}(t) c^{(\alpha)}_{\mathrm{d}}(t)\,, \tag{S26a}$$

$$0 = k_{\mathrm{on,s}} M^{(\alpha)}_{\mathrm{a,f}}(t) c^{(\alpha)}_{\mathrm{d}}(t) - k_{\mathrm{off,s}} M^{(\alpha)}_{\mathrm{a,b}}(t) \quad \Rightarrow \quad M^{(\alpha)}_{\mathrm{a,b}}(t) = K_{\mathrm{s}}\, M^{(\alpha)}_{\mathrm{a,f}}(t) c^{(\alpha)}_{\mathrm{d}}(t)\,, \tag{S26b}$$

$$0 = k_{\mathrm{on,m}} M^{(\alpha)}_{\mathrm{m,f}}(t) c^{(\alpha)}_{\mathrm{d}}(t) - k_{\mathrm{off,m}} M^{(\alpha)}_{\mathrm{m,b}}(t) \quad \Rightarrow \quad M^{(\alpha)}_{\mathrm{m,b}}(t) = K_{\mathrm{m}}\, M^{(\alpha)}_{\mathrm{m,f}}(t) c^{(\alpha)}_{\mathrm{d}}(t)\,, \tag{S26c}$$

where the equilibrium binding constants for inhibitor binding to monomers, fibril ends or fibril surfaces, given as:

$$K_{\mathrm{m}} = \frac{k_{\mathrm{on,m}}}{k_{\mathrm{off,m}}}\,, \quad K_{\mathrm{e}} = \frac{k_{\mathrm{on,e}}}{k_{\mathrm{off,e}}}\,, \quad K_{\mathrm{s}} = \frac{k_{\mathrm{on,s}}}{k_{\mathrm{off,s}}}\,. \tag{S27}$$

Defining the total concentration and mass of aggregates and monomers yields:

$$c^{(\alpha)}_{\mathrm{a}}(t) = c^{(\alpha)}_{\mathrm{a,f}}(t) + c^{(\alpha)}_{\mathrm{a,b}}(t) \quad \Rightarrow \quad c^{(\alpha)}_{\mathrm{a,f}}(t) = \frac{c^{(\alpha)}_{\mathrm{a}}(t)}{1 + K_{\mathrm{e}}\, c^{(\alpha)}_{\mathrm{d}}}\,, \tag{S28a}$$

$$M^{(\alpha)}_{\mathrm{a}}(t) = M^{(\alpha)}_{\mathrm{a,f}}(t) + M^{(\alpha)}_{\mathrm{a,b}}(t) \quad \Rightarrow \quad M^{(\alpha)}_{\mathrm{a,f}}(t) = \frac{M^{(\alpha)}_{\mathrm{a}}(t)}{1 + K_{\mathrm{s}}\, c^{(\alpha)}_{\mathrm{d}}}\,, \tag{S28b}$$

$$M^{(\alpha)}_{\mathrm{m}}(t) = M^{(\alpha)}_{\mathrm{m,f}}(t) + M^{(\alpha)}_{\mathrm{m,b}}(t) \quad \Rightarrow \quad M^{(\alpha)}_{\mathrm{m,f}}(t) = \frac{M^{(\alpha)}_{\mathrm{m}}(t)}{1 + K_{\mathrm{m}}\, c^{(\alpha)}_{\mathrm{d}}}\,. \tag{S28c}$$

In the limit of fast inhibitor binding to the target, we therefore arrive at the following system of kinetic equations

$$\frac{\mathrm{d}c_{\mathrm{a}}^{(\alpha)}}{\mathrm{d}t} = k_1\big(c_{\mathrm{d}}^{(\alpha)}\big)\big[M_{\mathrm{m}}^{(\alpha)}\big]^{n_1} + k_2\big(c_{\mathrm{d}}^{(\alpha)}\big)\big[M_{\mathrm{m}}^{(\alpha)}\big]^{n_2} M_{\mathrm{a}}^{(\alpha)}\,, \tag{S29a}$$

$$\frac{\mathrm{d}M_{\mathrm{a}}^{(\alpha)}}{\mathrm{d}t} = 2k_+\big(c_{\mathrm{d}}^{(\alpha)}\big)M_{\mathrm{m}}^{(\alpha)}\,c_{\mathrm{a}}^{(\alpha)}\,, \tag{S29b}$$

$$\frac{\mathrm{d}M_{\mathrm{m}}^{(\alpha)}}{\mathrm{d}t} = -2k_+\big(c_{\mathrm{d}}^{(\alpha)}\big)M_{\mathrm{m}}^{(\alpha)}\,c_{\mathrm{a}}^{(\alpha)} + \frac{J_{\mathrm{m}}^{(\alpha)}}{V^{(\alpha)}}\,, \tag{S29c}$$

$$\frac{\mathrm{d}c_{\mathrm{d}}^{(\alpha)}}{\mathrm{d}t} = \frac{J_{\mathrm{d}}^{(\alpha)}}{V^{(\alpha)}}\,, \tag{S29d}$$

where

$$k_1\big(c_{\mathrm{d}}^{(\alpha)}\big) = k_1\left(\frac{1}{1+K_{\mathrm{m}}c_{\mathrm{d}}^{(\alpha)}}\right)^{n_1}\,, \tag{S29e}$$

$$k_2\big(c_{\mathrm{d}}^{(\alpha)}\big) = k_2\left(\frac{1}{1+K_{\mathrm{m}}c_{\mathrm{d}}^{(\alpha)}}\right)^{n_2}\left(\frac{1}{1+K_{\mathrm{s}}c_{\mathrm{d}}^{(\alpha)}}\right)\,, \tag{S29f}$$

$$k_+\big(c_{\mathrm{d}}^{(\alpha)}\big) = k_+\left(\frac{1}{1+K_{\mathrm{m}}c_{\mathrm{d}}^{(\alpha)}}\right)\left(\frac{1}{1+K_{\mathrm{e}}c_{\mathrm{d}}^{(\alpha)}}\right)\,, \tag{S29g}$$

and $K_i$ is the equilibrium binding constant of the drug to monomers ($i=\mathrm{m}$), fibril ends ($i=\mathrm{e}$) and fibril surface sites ($i=\mathrm{s}$) and $k_1, k_2, k_+$ are the rate constants in the absence of the inhibitor. Eq. (S29) is Eq. (1) of the main text.

**S2.3. Solution to aggregation kinetics in compartment I.** By combining Eq. (S23) with Eq. (S29), we obtain the following "renormalized" kinetic equations for compartment I

$$\frac{\mathrm{d}c_{\mathrm{a}}^{\mathrm{I}}(t)}{\mathrm{d}t} = k_1\left(\frac{M_{\mathrm{m}}^{\mathrm{I}}(t)}{1+K_{\mathrm{m}}\,c_{\mathrm{d}}^{\mathrm{I}}}\right)^{n_1} + k_2\left(\frac{M_{\mathrm{m}}^{\mathrm{I}}(t)}{1+K_{\mathrm{m}}\,c_{\mathrm{d}}^{\mathrm{I}}}\right)^{n_2}\left(\frac{M_{\mathrm{m}}^{\mathrm{I}}(t)}{1+K_{\mathrm{s}}\,c_{\mathrm{d}}^{\mathrm{I}}}\right)\,, \tag{S30a}$$

$$\frac{\mathrm{d}M_{\mathrm{a}}^{\mathrm{I}}(t)}{\mathrm{d}t} = 2k_+\left(\frac{M_{\mathrm{m}}^{\mathrm{I}}(t)}{1+K_{\mathrm{m}}\,c_{\mathrm{d}}^{\mathrm{I}}}\right)\left(\frac{c_{\mathrm{a}}^{\mathrm{I}}(t)}{1+K_{\mathrm{e}}\,c_{\mathrm{d}}^{\mathrm{I}}}\right) = -\frac{\mathrm{d}M_{\mathrm{m}}^{\mathrm{I}}(t)}{\mathrm{d}t}\,, \tag{S30b}$$

with $M_{\mathrm{m}}^{\mathrm{I}}(0) = \xi_{\mathrm{m}}\Gamma_{\mathrm{m}}M_{\mathrm{m}}^{\mathrm{tot}}$ and $c_{\mathrm{d}}^{\mathrm{I}} = \xi_{\mathrm{d}}\Gamma_{\mathrm{d}}c_{\mathrm{d}}$. Conveniently, Eq. (S30) is equivalent to the kinetic equations in a homogeneous system Eq. (S1) but with effective rate parameters renormalized by the monomer and drug partitioning. The solution for the aggregate mass in compartment (I) is therefore given by (see Eq. (S3)):

$$\boxed{\frac{M_{\mathrm{m}}^{\mathrm{I}}(t)}{M_{\mathrm{m}}^{\mathrm{I}}(0)} = \left[1+\frac{\lambda_{\mathrm{I}}^2}{2\kappa_{\mathrm{I}}^2\theta}e^{\kappa_{\mathrm{I}}t}\right]^{-\theta}} \tag{S31}$$

where $\kappa_0 = \sqrt{2k_+k_2[M_{\mathrm{m}}^{\mathrm{tot}}]^{n_2+1}}$, $\lambda_0 = \sqrt{2k_+k_1[M_{\mathrm{m}}^{\mathrm{tot}}]^{n_1}}$, $\theta = \sqrt{2/[n_2(n_2+1)]}$ and

$$\kappa_{\mathrm{I}} = \kappa_0\;(\xi_{\mathrm{m}}\Gamma_{\mathrm{m}})^{\frac{n_2+1}{2}}\left(\frac{1}{1+K_{\mathrm{m}}\,\xi_{\mathrm{d}}\Gamma_{\mathrm{d}}c_{\mathrm{d}}}\right)^{\frac{n_2+1}{2}}\left(\frac{1}{1+K_{\mathrm{e}}\,\xi_{\mathrm{d}}\Gamma_{\mathrm{d}}c_{\mathrm{d}}}\right)^{\frac{1}{2}}\left(\frac{1}{1+K_{\mathrm{s}}\,\xi_{\mathrm{d}}\Gamma_{\mathrm{d}}c_{\mathrm{d}}}\right)^{\frac{1}{2}}\,, \tag{S32a}$$

$$\lambda_{\mathrm{I}} = \lambda_0\;(\xi_{\mathrm{m}}\Gamma_{\mathrm{m}})^{\frac{n_1}{2}}\left(\frac{1}{1+K_{\mathrm{m}}\,\xi_{\mathrm{d}}\Gamma_{\mathrm{d}}c_{\mathrm{d}}}\right)^{\frac{n_1+1}{2}}\left(\frac{1}{1+K_{\mathrm{e}}\,\xi_{\mathrm{d}}\Gamma_{\mathrm{d}}c_{\mathrm{d}}}\right)^{\frac{1}{2}}\,. \tag{S32b}$$

Similarly, the solution for the aggregate number concentration in compartment I is

$$\boxed{\frac{c_{\mathrm{a}}^{\mathrm{I}}(t)}{c_{\mathrm{a}}^{\mathrm{I}}(\infty)} = \left[1+\frac{2\kappa_{\mathrm{I}}^2\theta}{\lambda_{\mathrm{I}}^2}e^{-\kappa_{\mathrm{I}}t}\right]^{-1}} \tag{S33}$$

where the asymptotic aggregate concentration is

$$c_{\mathrm{a}}^{\mathrm{I}}(\infty) = \frac{1}{2k_+}\sqrt{\kappa_{\mathrm{I}}^2\theta^2+\frac{2\lambda_{\mathrm{I}}^2}{n_1\theta}} \simeq \frac{\kappa_{\mathrm{I}}\theta}{2k_+}, \tag{S34}$$

i.e.

$$c_{\mathrm{a}}^{\mathrm{I}}(\infty) \simeq \frac{\kappa_0\theta}{2k_+}(\xi_{\mathrm{m}}\Gamma_{\mathrm{m}})^{\frac{n_2+1}{2}}\left(\frac{1}{1+K_{\mathrm{m}}\,\xi_{\mathrm{d}}\Gamma_{\mathrm{d}}c_{\mathrm{d}}}\right)^{\frac{n_2+1}{2}}\left(\frac{1}{1+K_{\mathrm{e}}\,\xi_{\mathrm{d}}\Gamma_{\mathrm{d}}c_{\mathrm{d}}}\right)^{-\frac{1}{2}}\left(\frac{1}{1+K_{\mathrm{s}}\,\xi_{\mathrm{d}}\Gamma_{\mathrm{d}}c_{\mathrm{d}}}\right)^{\frac{1}{2}}. \quad \text{[S35]}$$

The performance of Eq. (S31) and Eq. (S33) against numerical integration of Eq. (S15) is shown in Figs. S2 and S3 for the cases when the inhibitor binds monomers or fibril surface sites.

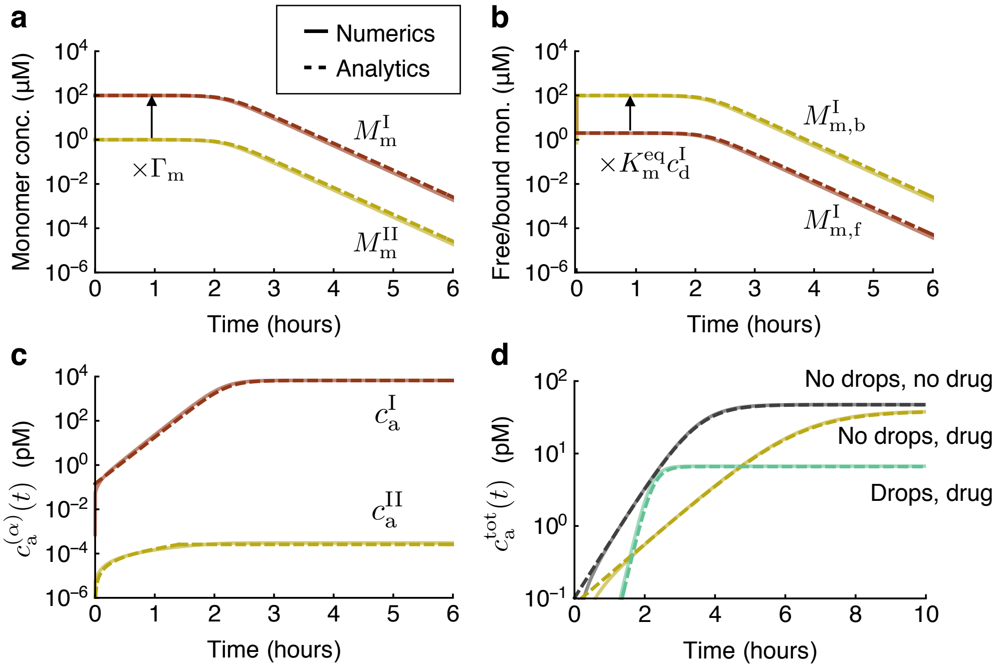

**Fig. S2. Comparison between numerical (solid lines) and analytical (dashed lines) solution to (S4) for a monomer binder.** **(a)** Monomer concentrations $M_{\mathrm{m}}^{(\alpha)}(t)$ inside ($\alpha$ = I) and outside ($\alpha$ = II) of the droplet. Solid lines are the numerical solution, while dashed lines are Eq. (S31) and Eq. (S36). Note that $M_{\mathrm{m}}^{\mathrm{I}}(t) = \Gamma_{\mathrm{m}} M_{\mathrm{m}}^{\mathrm{II}}(t)$ holds at all times. **(b)** Concentrations of free and bound monomers inside compartment I. Dashed lines are obtained using Eq. (S28). **(c)** Aggregate concentrations $c_{\mathrm{a}}^{(\alpha)}(t)$ inside ($\alpha$ = I) and outside ($\alpha$ = II) of the droplet in the presence of a drug that binds surface ends. Solid lines are from the numerical solution of Eq. (S4). Dashed lines are Eq. (S33) respectively Eq. (S37). **(d)** Total aggregate concentration in the absence of drops and drug (black), in the absence of drops but in the presence of drugs (yellow), and in the presence of drops and drug (green). Parameters are: $k_+ = 3\times10^6$ M$^{-1}$s$^{-1}$, $k_1 = 10^{-4}$ M$^{-1}$s$^{-1}$, $k_2 = 4\times10^4$ M$^{-2}$s$^{-1}$, $n_1 = n_2 = 2$, $M_{\mathrm{m}}^{\mathrm{tot}} = 1\,\mu$M, $c_{\mathrm{d}} = 1\,\mu$M, $k_{\mathrm{on,s}} = 10^5$ M$^{-1}$s$^{-1}$, $k_{\mathrm{off,s}} = 2\times10^{-1}$ s$^{-1}$, $\Gamma_{\mathrm{d}} = \Gamma_{\mathrm{m}} = 100$, $V^{\mathrm{I}}/V = 10^{-3}$. $k$ is set to a large number (absolute value does not matter for kinetics). The chosen values for the rate constants for aggregation correspond to typical values measured for A$\beta$42 aggregation in vitro (8).

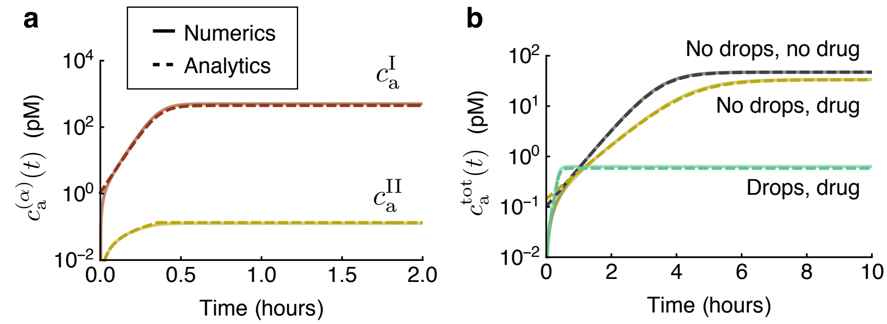

**Fig. S3. Comparison between numerical (solid lines) and analytical (dashed lines) solution to (S4) for an inhibitor that binds fibril surface sites.** **(a)** Aggregate concentrations $c_{\mathrm{a}}^{(\alpha)}(t)$ inside ($\alpha$ = I) and outside ($\alpha$ = II) of the droplet in the presence of a drug that binds surface ends. Solid lines are from the numerical solution of Eq. (S4). Dashed lines are Eq. (S33) respectively Eq. (S37). **(b)** Total aggregate concentration in the absence of drops and drug (black), in the absence of drops but in the presence of drugs (yellow), and in the presence of drops and drug (green). Parameters are: $k_+ = 3\times10^6$ M$^{-1}$s$^{-1}$, $k_1 = 10^{-4}$ M$^{-1}$s$^{-1}$, $k_2 = 4\times10^4$ M$^{-2}$s$^{-1}$, $n_1 = n_2 = 2$, $M_{\mathrm{m}}^{\mathrm{tot}} = 1\,\mu$M, $c_{\mathrm{d}} = 1\,\mu$M, $k_{\mathrm{on,s}} = 10^5$ M$^{-1}$s$^{-1}$, $k_{\mathrm{off,s}} = 10^{-1}$ s$^{-1}$, $\Gamma_{\mathrm{d}} = \Gamma_{\mathrm{m}} = 10$, $V^{\mathrm{I}}/V = 10^{-3}$. The chosen values for the rate constants for aggregation correspond to typical values measured for A$\beta$42 aggregation in vitro (8).

**S2.4. Solution to aggregation kinetics in compartment II.** The concentration of monomers in compartment II is obtained from Eq. (S31) using Eq. (S19), yielding:

$$\boxed{\frac{M_{\mathrm{m}}^{\mathrm{II}}(t)}{M_{\mathrm{m}}^{\mathrm{II}}(0)} = \left[1 + \frac{\lambda_{\mathrm{I}}^2}{2\kappa_{\mathrm{I}}^2\theta} e^{\kappa_{\mathrm{I}} t}\right]^{-\theta}} \tag{S36}$$

Note that, due to Eq. (S19), the dynamics of $M_{\mathrm{m}}^{\mathrm{II}}(t)$ is entirely determined by $M_{\mathrm{m}}^{\mathrm{I}}(t)$. This conclusion is verified against numerical integration of Eq. (S15) in Fig. S2(a).

To obtain an expression for the aggregate number concentration in compartment II, we note that, for strong monomer partitioning into compartment I ($\Gamma_{\mathrm{m}} \gg 1$), aggregation kinetics is much faster inside compartment (I) compared to compartment (II). As a result, aggregation inside compartment (I) reaches the plateau while aggregation inside compartment (II) is still in the exponential growth phase. The solution for the aggregate number concentration inside compartment (II) is thus described by the exponential function:

$$c_{\mathrm{a}}^{\mathrm{II}}(t) \simeq \frac{\nu}{\kappa_{\mathrm{II}}} \sinh\left(\kappa_{\mathrm{II}} t\right) , \tag{S37}$$

where

$$\nu = k_1 \left(M_{\mathrm{m}}^{\mathrm{tot}}\right)^{n_1} \xi_{\mathrm{m}}^{n_1} \left(\frac{1}{1 + K_{\mathrm{m}}\, \xi_{\mathrm{d}} \Gamma_{\mathrm{d}} c_{\mathrm{d}}}\right)^{n_1} \tag{S38}$$

and

$$\kappa_{\mathrm{II}} = \kappa_0\, \xi_{\mathrm{m}}^{\frac{n_2+1}{2}} \left(\frac{1}{1 + K_{\mathrm{m}}\, \xi_{\mathrm{d}} c_{\mathrm{d}}}\right)^{\frac{n_2+1}{2}} \left(\frac{1}{1 + K_{\mathrm{e}}\, \xi_{\mathrm{d}} c_{\mathrm{d}}}\right)^{\frac{1}{2}} \left(\frac{1}{1 + K_{\mathrm{s}}\, \xi_{\mathrm{d}} c_{\mathrm{d}}}\right)^{\frac{1}{2}} . \tag{S39}$$

When aggregation in compartment (I) reaches the plateau, aggregation inside compartment (II) must stop, due to mass conservation. An estimate for this stopping time is the half-time of aggregation inside compartment (I):

$$t_{1/2} \simeq \frac{1}{\kappa_{\mathrm{I}}} \ln\left(\frac{2\kappa_{\mathrm{I}}^2\theta}{\lambda_{\mathrm{I}}^2}\right) \propto \frac{1}{\kappa_{\mathrm{I}}}. \tag{S40}$$

Thus, aggregation inside compartment (II) will plateau at the value

$$c_{\mathrm{a}}^{\mathrm{II}}(\infty) \simeq \frac{\nu}{\kappa_{\mathrm{II}}} \sinh\left[\frac{\kappa_{\mathrm{II}}}{\kappa_{\mathrm{I}}} \ln\left(\frac{2\kappa_{\mathrm{I}}^2\theta}{\lambda_{\mathrm{I}}^2}\right)\right] . \tag{S41}$$

In summary, the approximate solution describing the time course of the aggregate number concentration in compartment II is given by

$$\boxed{c_{\mathrm{a}}^{\mathrm{II}}(t) \simeq \begin{cases} \frac{\nu}{\kappa_{\mathrm{II}}} \sinh\left(\kappa_{\mathrm{II}} t\right) , & t < \frac{1}{\kappa_{\mathrm{I}}} \ln\left(\frac{2\kappa_{\mathrm{I}}^2\theta}{\lambda_{\mathrm{I}}^2}\right) \\ \frac{\nu}{\kappa_{\mathrm{II}}} \sinh\left[\frac{\kappa_{\mathrm{II}}}{\kappa_{\mathrm{I}}} \ln\left(\frac{2\kappa_{\mathrm{I}}^2\theta}{\lambda_{\mathrm{I}}^2}\right)\right] , & t > \frac{1}{\kappa_{\mathrm{I}}} \ln\left(\frac{2\kappa_{\mathrm{I}}^2\theta}{\lambda_{\mathrm{I}}^2}\right) \end{cases}} \tag{S42}$$

We can simplify the above expressions further by expanding the sinh function in Eq. (S41) to leading order for $\Gamma_{\mathrm{m}} \gg 1$ ($\sinh(x) \simeq x + \cdots$, $x \ll 1$),* yielding

$$c_{\mathrm{a}}^{\mathrm{II}}(\infty) \simeq \frac{\nu}{\kappa_{\mathrm{I}}} \ln\left(\frac{2\kappa_{\mathrm{I}}^2\theta}{\lambda_{\mathrm{I}}^2}\right) . \tag{S43}$$

Neglecting the logarithmic contribution in front of the power law dependence on $\kappa_{\mathrm{I}}$, $c_{\mathrm{a}}^{\mathrm{II}}(\infty)$ is given approximately by

$$c_{\mathrm{a}}^{\mathrm{II}}(\infty) \simeq \frac{\nu}{\kappa_{\mathrm{I}}} , \tag{S44}$$

which can be written explicitly as

$$c_{\mathrm{a}}^{\mathrm{II}}(\infty) \simeq w\, \frac{\kappa_0\theta}{2k_+}\, \xi_{\mathrm{m}}^{n_1 - \frac{n_2+1}{2}} \Gamma_{\mathrm{m}}^{-\frac{n_2+1}{2}} \left(\frac{1}{1 + K_{\mathrm{m}}\, \xi_{\mathrm{d}} \Gamma_{\mathrm{d}} c_{\mathrm{d}}}\right)^{n_1 - \frac{n_2+1}{2}} \left(\frac{1}{1 + K_{\mathrm{e}}\, \xi_{\mathrm{d}} \Gamma_{\mathrm{d}} c_{\mathrm{d}}}\right)^{-\frac{1}{2}} \left(\frac{1}{1 + K_{\mathrm{s}}\, \xi_{\mathrm{d}} \Gamma_{\mathrm{d}} c_{\mathrm{d}}}\right)^{-\frac{1}{2}} , \tag{S45}$$

where

$$w = \frac{k_1 \left(M_{\mathrm{m}}^{\mathrm{tot}}\right)^{n_1 - n_2 - 1}}{k_2\theta} \ll 1. \tag{S46}$$

*Note that $\frac{\kappa_{\mathrm{II}}}{\kappa_{\mathrm{I}}} \simeq \Gamma_{\mathrm{m}}^{-\frac{n_2+1}{2}} \ll 1$ for $\Gamma_{\mathrm{m}} \gg 1$.

**S2.5. Summary.** In summary, we have obtained the following asymptotic expressions for the terminal ($t \to \infty$) aggregate concentration in the different scenarios:

- **1) No drops, no drug.**

$$c_{\mathrm{a}}(\infty)|_{\mathrm{homo},c_{\mathrm{d}}=0} \simeq \frac{\kappa_0 \theta}{2k_+}. \qquad \text{[S47]}$$

- **2) No drops + drug.**

$$c_{\mathrm{a}}(\infty)|_{\mathrm{homo}} \simeq \frac{\kappa_0 \theta}{2k_+} \left(\frac{1}{1+K_{\mathrm{m}}\, c_{\mathrm{d}}}\right)^{\frac{n_2+1}{2}} \left(\frac{1}{1+K_{\mathrm{e}}\, c_{\mathrm{d}}}\right)^{-\frac{1}{2}} \left(\frac{1}{1+K_{\mathrm{s}}\, c_{\mathrm{d}}}\right)^{\frac{1}{2}}. \qquad \text{[S48]}$$

- **3) Drops, no drug.**

$$c_{\mathrm{a}}^{\mathrm{I}}(\infty)\big|_{c_{\mathrm{d}}=0} \simeq \frac{\kappa_0 \theta}{2k_+} (\xi_{\mathrm{m}} \Gamma_{\mathrm{m}})^{\frac{n_2+1}{2}}, \qquad \text{[S49]}$$

$$c_{\mathrm{a}}^{\mathrm{II}}(\infty)\big|_{c_{\mathrm{d}}=0} \simeq w\, \frac{\kappa_0 \theta}{2k_+}\, \xi_{\mathrm{m}}^{n_1 - \frac{n_2+1}{2}} \Gamma_{\mathrm{m}}^{-\frac{n_2+1}{2}}. \qquad \text{[S50]}$$

- **4) Drops + drug.**

$$c_{\mathrm{a}}^{\mathrm{I}}(\infty) \simeq \frac{\kappa_0 \theta}{2k_+} (\xi_{\mathrm{m}} \Gamma_{\mathrm{m}})^{\frac{n_2+1}{2}} \left(\frac{1}{1+K_{\mathrm{m}}\, \xi_{\mathrm{d}} \Gamma_{\mathrm{d}} c_{\mathrm{d}}}\right)^{\frac{n_2+1}{2}} \left(\frac{1}{1+K_{\mathrm{e}}\, \xi_{\mathrm{d}} \Gamma_{\mathrm{d}} c_{\mathrm{d}}}\right)^{-\frac{1}{2}} \left(\frac{1}{1+K_{\mathrm{s}}\, \xi_{\mathrm{d}} \Gamma_{\mathrm{d}} c_{\mathrm{d}}}\right)^{\frac{1}{2}}, \qquad \text{[S51]}$$

$$c_{\mathrm{a}}^{\mathrm{II}}(\infty) \simeq w\, \frac{\kappa_0 \theta}{2k_+}\, \xi_{\mathrm{m}}^{n_1 - \frac{n_2+1}{2}} \Gamma_{\mathrm{m}}^{-\frac{n_2+1}{2}} \left(\frac{1}{1+K_{\mathrm{m}}\, \xi_{\mathrm{d}} \Gamma_{\mathrm{d}} c_{\mathrm{d}}}\right)^{n_1 - \frac{n_2+1}{2}} \left(\frac{1}{1+K_{\mathrm{e}}\, \xi_{\mathrm{d}} \Gamma_{\mathrm{d}} c_{\mathrm{d}}}\right)^{-\frac{1}{2}} \left(\frac{1}{1+K_{\mathrm{s}}\, \xi_{\mathrm{d}} \Gamma_{\mathrm{d}} c_{\mathrm{d}}}\right)^{-\frac{1}{2}}. \qquad \text{[S52]}$$

### S3. Calculation of enhancement function $\mathcal{E}$

As discussed in the main text, we define the enhancement function $\mathcal{E}$ as:

$$\mathcal{E} = \frac{c_{\mathrm{a}}(\infty)|_{\mathrm{homo}}}{\bar{c_{\mathrm{a}}}(\infty)}, \qquad \text{[S53]}$$

where the average concentration of aggregates formed in the presence of a liquid drop is calculated as:

$$\bar{c_{\mathrm{a}}}(\infty) = \frac{V^{\mathrm{I}} c_{\mathrm{a}}^{\mathrm{I}}(\infty) + V^{\mathrm{II}} c_{\mathrm{a}}^{\mathrm{II}}(\infty)}{V}, \qquad \text{[S54]}$$

with $c_{\mathrm{a}}^{\mathrm{I}}(\infty)$ and $c_{\mathrm{a}}^{\mathrm{II}}(\infty)$ given by Eq. (S51) and Eq. (S52). The asymptotic aggregate number concentration in the presence of a drug but without compartments is given by Eq. (S48).

Using Eq. (S49) and Eq. (S51), we find the following leading contribution to the asymptotic concentration in I compared to the uninhibited case:

$$\frac{c_{\mathrm{a}}^{\mathrm{I}}(\infty)}{c_{\mathrm{a}}^{\mathrm{I}}(\infty)\big|_{c_{\mathrm{d}}=0}} \simeq (\xi_{\mathrm{m}} \Gamma_{\mathrm{m}})^{\frac{n_2+1}{2}} \left(\frac{1+K_\times c_{\mathrm{d}}}{1+\xi_{\mathrm{d}} \Gamma_{\mathrm{d}}\, K_\times c_{\mathrm{d}}}\right)^{\beta_1} \quad \text{with} \quad \beta_1 = \begin{cases} \frac{n_2+1}{2} & \text{(monomers)} \\ -\frac{1}{2} & \text{(ends)} \\ \frac{1}{2} & \text{(surface)} \end{cases} \qquad \text{[S55]}$$

and $\times =$ m,e,s.. Similarly, using Eq. (S50) and Eq. (S52) we can write the asymptotic concentration in compartment II compared to the uninhibited systemas:

$$\frac{c_{\mathrm{a}}^{\mathrm{II}}(\infty)}{c_{\mathrm{a}}^{\mathrm{II}}(\infty)\big|_{c_{\mathrm{d}}=0}} \simeq w\, \xi_{\mathrm{m}}^{n_1 - \frac{n_2+1}{2}} \Gamma_{\mathrm{m}}^{-\frac{n_2+1}{2}} (1+K_\times c_{\mathrm{d}})^{\beta_1} (1+\xi_{\mathrm{d}} \Gamma_{\mathrm{d}}\, K_\times c_{\mathrm{d}})^{\beta_2} \quad \text{with} \quad \beta_2 = \begin{cases} \frac{n_2+1}{2} - n_1, & \text{(monomers)} \\ \frac{1}{2}, & \text{(ends)} \\ \frac{1}{2} & \text{(surface)} \end{cases}. \qquad \text{[S56]}$$

Eq. (S55) and Eq. (S56) yield Eqs. (2) and (3) of the main text.

### S4. Potency increase

We now study the role of liquid condensates on drug potency. As discussed in the main text, we define the drug-response curve $\mathcal{R}(c_\mathrm{d})$ as in the main text as the relative difference between asymptotic aggregate concentrations obtained in the presence of droplets and drug and the homogeneous system

$$\mathcal{R}(c_\mathrm{d}) = 1 - \frac{\bar{c_\mathrm{a}}(\infty)}{\bar{c_\mathrm{a}}(\infty)|_{c_\mathrm{d}=0}}. \qquad \text{[S57]}$$

The potency refers to the concentration $\mathrm{EC}_{50}$ of a drug necessary to reach 50% of the maximal effect characterized by the drug-response, i.e. $\mathcal{R}(\mathrm{EC}_{50}) = 1/2$. A drug is said to be more potent if a lower drug concentration required to reach half of the maximal response, i.e., half of the efficacy. We define

$$\mathcal{P} = \frac{1}{\mathrm{EC}_{50}}. \qquad \text{[S58]}$$

**Homogeneous system.** In this case, we use Eq. (S47) and Eq. (S48)

$$c_\mathrm{a}(\infty)|_{\mathrm{homo},c_\mathrm{d}=0} \simeq \frac{\kappa_0\theta}{2k_+}, \quad c_\mathrm{a}(\infty)|_{\mathrm{homo}} \simeq \frac{\kappa_0\theta}{2k_+}\left(\frac{1}{1+K_\times c_\mathrm{d}}\right)^{\beta_1} \qquad \text{[S59]}$$

to compute the response function as

$$\mathcal{R} = 1 - \left(\frac{1}{1+K_\times c_\mathrm{d}}\right)^{\beta_1}. \qquad \text{[S60]}$$

Thus, the concentration of the drug that is required to yield half of its efficacy is given by

$$K_\times c_\mathrm{d} = 2^{1/\beta_1} - 1 \quad \Rightarrow \quad \mathrm{EC}_{50}|_{\mathrm{homo}} = \frac{2^{1/\beta_1}-1}{K_\times} \quad \Rightarrow \quad \mathcal{P}_\times|_{\mathrm{homo}} = \frac{1}{\mathrm{EC}_{50}|_{\mathrm{homo}}} = \frac{K_\times}{2^{1/\beta_1}-1}. \qquad \text{[S61]}$$

**Phase separated system.** In the presence of a liquid compartment, we have from Eq. (S49), Eq. (S50), Eq. (S51) and Eq. (S52)

$$\bar{c_\mathrm{a}}(\infty)|_{c_\mathrm{d}=0} \simeq \frac{V^\mathrm{I}}{V}\frac{\kappa_0\theta}{2k_+}(\xi_\mathrm{m}\Gamma_\mathrm{m})^{\frac{n_2+1}{2}} + w\frac{V^\mathrm{II}}{V}\frac{\kappa_0\theta}{2k_+}\xi_\mathrm{m}^{n_1-\frac{n_2+1}{2}}\Gamma_\mathrm{m}^{-\frac{n_2+1}{2}} \qquad \text{[S62]}$$

and

$$\bar{c_\mathrm{a}}(\infty) \simeq \frac{V^\mathrm{I}}{V}\frac{\kappa_0\theta}{2k_+}(\xi_\mathrm{m}\Gamma_\mathrm{m})^{\frac{n_2+1}{2}}\left(\frac{1}{1+\xi_\mathrm{d}\Gamma_\mathrm{d}K_\times c_\mathrm{d}}\right)^{\beta_1} + w\frac{V^\mathrm{II}}{V}\frac{\kappa_0\theta}{2k_+}\xi_\mathrm{m}^{n_1-\frac{n_2+1}{2}}\Gamma_\mathrm{m}^{-\frac{n_2+1}{2}}(1+\xi_\mathrm{d}\Gamma_\mathrm{d}K_\times c_\mathrm{d})^{\beta_2}. \qquad \text{[S63]}$$

For $\Gamma_\mathrm{m} \gg 1$ and in the regime when the drop with drug system is the optimal one for suppressing aggregation (see phase diagrams in Fig. 3 of the main text), we can neglect the terms proportional to $w$ in the two equations above, yielding

$$\mathcal{R} \simeq 1 - \left(\frac{1}{1+\xi_\mathrm{d}\Gamma_\mathrm{d}K_\times c_\mathrm{d}}\right)^{\beta_1}. \qquad \text{[S64]}$$

Thus,

$$\xi_\mathrm{d}\Gamma_\mathrm{d}K_\times c_\mathrm{d} = 2^{1/\beta_1}-1 \quad \Rightarrow \quad \mathrm{EC}_{50,\mathrm{drops}} = \frac{2^{1/\beta_1}-1}{K_\times\xi_\mathrm{d}\Gamma_\mathrm{d}} \quad \Rightarrow \quad \mathcal{P}_\times|_{\mathrm{drops}} = \frac{1}{\mathrm{EC}_{50,\mathrm{drops}}} = \frac{K_\times\xi_\mathrm{d}\Gamma_\mathrm{d}}{2^{1/\beta_1}-1}. \qquad \text{[S65]}$$

**Relative potency.** Combining Eq. (S61) and Eq. (S65), we obtain the universal, i.e., inhibition mechanism-independent, relative potency:

$$\mathcal{P} = \frac{\mathrm{EC}_{50}|_{\mathrm{homo}}}{\mathrm{EC}_{50}|_{\mathrm{drops}}} \simeq \xi_\mathrm{d}\Gamma_\mathrm{d} = \frac{\Gamma_\mathrm{d}}{1+(\Gamma_\mathrm{d}-1)V^\mathrm{I}/V}, \qquad \text{[S66]}$$

which is Eq. (4) of the main text.

### S5. Glossary

**Table S2.** List of symbols.

| Symbol | Meaning |
|---|---|
| $c_\mathrm{a}(t)$ | aggregate number concentration |
| $c_\mathrm{a}(\infty)$ | terminal (i.e. $t \to \infty$) aggregate number concentration |
| $M_\mathrm{a}(t)$ | aggregate mass concentration |
| $M_\mathrm{m}(t)$ | monomer concentration |
| $M_\mathrm{m}^\mathrm{tot}$ | total concentration of monomer in system (conserved) |
| $k_1$ | rate constant for primary nucleation |
| $k_2$ | rate constant for secondary nucleation |
| $k_+$ | rate constant for aggregate elongation (growth) |
| $n_1$ | reaction order for primary nucleation |
| $n_2$ | reaction order for secondary nucleation |
| $c_\mathrm{a,f}(t)$ | free aggregate number concentration |
| $c_\mathrm{a,b}(t)$ | bound aggregate number concentration |
| $M_\mathrm{a,f}(t)$ | free aggregate mass concentration |
| $M_\mathrm{a,b}(t)$ | bound aggregate mass concentration |
| $M_\mathrm{m,f}(t)$ | free monomer concentration |
| $M_\mathrm{m,b}(t)$ | bound monomer concentration |
| $c_\mathrm{d}(t)$ | (free) inhibitor concentration |
| $k_\mathrm{on,e}$ | on-rate constant for inhibitor binding to aggregate ends |
| $k_\mathrm{off,e}$ | off-rate constant for inhibitor binding to aggregate ends |
| $K_\mathrm{e}$ | $\frac{k_\mathrm{on,e}}{k_\mathrm{off,e}}$ binding constant for inhibitor binding to aggregate ends |
| $k_\mathrm{on,s}$ | on-rate constant for inhibitor binding to aggregate surface |
| $k_\mathrm{off,s}$ | off-rate constant for inhibitor binding to aggregate surface |
| $K_\mathrm{s}$ | $\frac{k_\mathrm{on,s}}{k_\mathrm{off,s}}$ equilibrium constant for inhibitor binding to aggregate surface |
| $k_\mathrm{on,m}$ | on-rate constant for inhibitor binding to monomers |
| $k_\mathrm{off,m}$ | off-rate constant for inhibitor binding to monomers |
| $K_\mathrm{m}$ | $\frac{k_\mathrm{on,m}}{k_\mathrm{off,m}}$ equilibrium constant for inhibitor binding to monomers |
| $\theta$ | $\sqrt{\frac{2}{n_2(n_2+1)}}$ |
| $\lambda_0$ | $\sqrt{2k_+ k_1 [M_\mathrm{m}^\mathrm{tot}]^{n_1}}$ rate of aggregate proliferation through primary nucleation and growth |
| $\kappa_0$ | $\sqrt{2k_+ k_2 [M_\mathrm{m}^\mathrm{tot}]^{n_2+1}}$ rate of aggregate proliferation through secondary nucleation and growth |
| $\Gamma_\mathrm{m}$ | monomer partition coefficient |
| $\Gamma_\mathrm{d}$ | drug partition coefficient |
| $\xi_\mathrm{m}$ | $\frac{1}{1+(\Gamma_\mathrm{m}-1)V^\mathrm{I}/V}$ partitioning degree of monomers |
| $\xi_\mathrm{d}$ | $\frac{1}{1+(\Gamma_\mathrm{d}-1)V^\mathrm{I}/V}$ partitioning degree of drug |
| $\alpha =$I,II | Compartment I or II |
| $V$ | System volume |
| $V^\mathrm{I}$ and $V^\mathrm{II}$ | Volume of compartment I or II |
| $J$ | Diffusive flux |
| $D$ | Diffusion coefficient |
| $\mathcal{E}$ | Enhancement function |
| $\mathcal{R}$ | Response function |
| $\mathcal{P}$ | Potency |
| $\mathcal{P}_\mathrm{rel}$ | Relative potency |
